# rRNA intermediates dictate nucleolar architecture in *C. elegans*

**DOI:** 10.1101/2025.08.07.669057

**Authors:** Jiewei Cheng, Liang Liu, Demin Xu, Yan Kuang, Ao Zheng, Minjie Hong, Xiaona Huang, Xiaotian Zhou, Wanjing Niu, Xinya Huang, Chengming Zhu, Xiangyang Chen, Xuezhu Feng, Shouhong Guang

## Abstract

The nucleolus is a multilayered, membraneless organelle essential for rRNA transcription, processing and ribosome assembly; however, it remains largely unclear whether and how these molecular events coordinate the organization of the multilayered nucleolar structure. We previously reported that the accumulation of 27SA_2_ rRNA upon the loss of a particular subset of RPL proteins elicits nucleolar reshaping, resulting in the formation of nucleolar rings and vacuoles in *C. elegans.* Here, we systematically investigated the effects of pre-18S rRNA processing and rRNA transcription on nucleolar structure and reported that defects in these molecular events result in reorganization of the nucleolar architecture. In particular, the depletion of a special subset of small ribosomal subunit proteins (RPSs), namely, class I RPSs, induces a concentric three-layered nucleolar architecture; the inhibition of rRNA transcription induces another three-layered architecture, a sandwich-like nucleoli. Via fluorescence imaging, we found that nucleolar proteins undergo demixing and spatial redistribution into distinct subnucleolar regions during nucleolar structure reorganization. Tracking the subcellular localization of rDNA via a lacI::tagRFP/lacO system revealed that rDNA also redistributed upon nucleolar reorganization. We further identified the Ce.22S rRNA intermediate, which accumulates upon the loss of class I RPSs and regulates the reorganization of the nucleolar structure. Moreover, the conserved nucleolar protein NUCL-1/nucleolin is essential for nucleolar architecture reorganization and regulates the development of nematodes during impaired rRNA production. Together, these findings reveal that the nucleolar structure is orchestrated by the status of rRNA biogenesis and that the accumulation of particular rRNA intermediates can direct the organization of specific nucleolar architectures and the spatial mixing/demixing behavior of the nucleolar proteome.

## Introduction

Eukaryotic cells achieve spatiotemporal regulation of multiple biochemical reactions through compartmentalized intracellular structures. Among the many subcellular structures, the nucleolus, a multilayered membraneless organelle in the nucleus, serves primarily for ribosomal RNA (rRNA) synthesis and ribosome assembly ^1–6^. In eukaryotic cells, two distinct types of nucleolar architectures have been identified, including the tripartite nucleolus and the bipartite nucleolus. The tripartite nucleolus, found in mammals and plants, consists of three internal compartments organized from the periphery to the center: the granular component (GC), the dense fibrillar component (DFC), and the fibrillar center (FC), each of which performs specialized functions in rRNA synthesis, rRNA processing and ribosome assembly, respectively ^7–10^. In contrast, other eukaryotes, such as *Drosophila* ^11^, possess a bipartite nucleolus, which lacks a distinct FC layer. Similarly, recent studies utilizing superresolution microscopy have revealed that the *C. elegans* nucleolus is also partitioned into two intermingled subcompartments, the FC and GC (Fig. 1A) ^12^.

**Figure 1.**
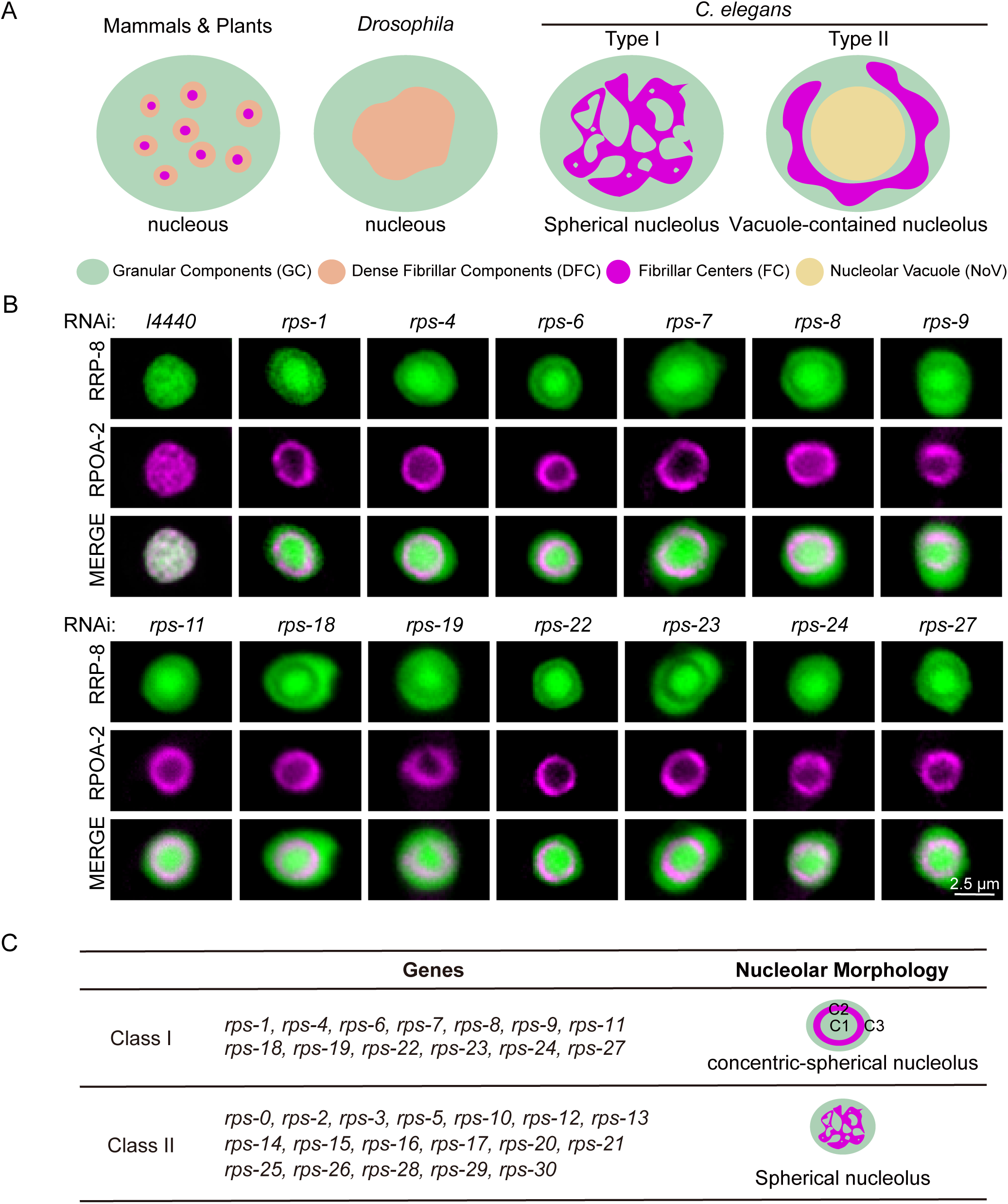
Knocking down class I *rps* genes induces nucleolar reorganization. (A) Schematic diagram of nucleolar organization in mammals, plants, *Drosophila* and *C. elegans*. (B) Fluorescence microscopy images of *C. elegans* nucleoli in hypodermal cells after the indicated genes were knocked down via RNAi. All worms were imaged at the L3–L4 stage. Scale bar, 2.5 μm. (C) Summary of concentric-spherical nucleoli formation upon the knockdown of class I and II *rps* genes.

Despite species-specific structural differences, the mechanisms of ribosome biogenesis are highly conserved among eukaryotes. Insights into ribosome assembly have largely been derived from yeast models, underscoring the universal importance of this process ^5,13,14^. The nucleolus originates from nucleolar organizer regions (NORs) ^2^, which consist of rDNA repeat clusters located primarily on chromosomes I and V in *C. elegans* ^15^. The core 47S rRNA is transcribed from rDNA by RNA polymerase I and appears as a characteristic “Christmas tree” structure under electron microscopy, reflecting active transcription. This precursor rRNA is subsequently processed into mature 18S, 5.8S, and 26S rRNAs ^16,17^. In mammals, the transcription of nascent pre-rRNA initiates at the FC-DFC interface and proceeds outward through the nucleolus. As it moves, the rRNA undergoes sequential processing and modification. Continuous production and maturation of rRNA are crucial for preserving both the architecture and the functional dynamics of the nucleolus ^18,19^. Mature ribosomes consist of two unequal subunits: the 40S small subunit (SSU) and the 60S large subunit (LSU). The 40S subunit comprises 18S rRNA and 33 small subunit ribosomal proteins (RPSs), whereas the 60S subunit contains 25S/28S rRNA, 5.8S rRNA, and 47 large subunit ribosomal proteins (RPLs) ^20–24^.

Ribosome biogenesis is intimately linked with nucleolar structure organization. Perturbations in nucleolar function often manifest as distinct morphological changes. For example, treatment with actinomycin D, an inhibitor of RNA polymerase I, causes the relocation of FC and DFC to the nucleolus periphery and the formation of structures known as nucleolar caps in mammalian cells ^25–27^. Nucleolar hypertrophy, which reflects the enhanced ribosomal production required to sustain rapid cell proliferation, has emerged as a hallmark of many tumors ^28,29^. In our previous study, knockdown of the class I *rpl* genes in *C. elegans* led to inappropriate accumulation of the 27SA_2_ rRNA intermediate, altered the nucleolar morphology, and resulted in the formation of nucleolar rings and nucleolar vacuoles (NoVs) ^3^. Nucleolar proteins, which are usually involved in rRNA transcription and processing, are excluded from NoVs; nevertheless, nucleoplasmic proteins accumulate in NoV.

To further investigate the regulation of nucleolar structure organization, we examined whether deficient RPS proteins and impaired rRNA production would also induce nucleolar reshaping or nucleolar architecture alteration in *C. elegans*. Unlike the formation of NoV, which is induced by class I *rpl* knockdown ^3^, we observed two striking nucleolar morphologies, concentric-spherical nucleoli and sandwich-like nucleoli. Concentric-spherical nucleoli are induced by the knockdown of 13 particular RPS proteins, which are termed class I RPSs. Interestingly, after the class I RPS proteins were knocked down, the Ce.22S rRNA intermediate consistently accumulated. Moreover, the proteins involved in rRNA transcription as well as 18S and 26S rRNA processing exhibit a demixing behavior and are separated into particular nucleolar subcompartments. The formation of sandwich-like nucleoli is induced by actinomycin D treatment, in which rRNA transcription as well as 18S and 26S rRNA processing factors also exhibit demixing behavior. We identified NUCL-1 as a key factor in nucleolar structure reorganization and as a checkpoint that promotes the development of *C. elegans* under conditions of impaired rRNA production and processing. These findings highlight the critical interplay between nucleolar architecture and rRNA biogenesis and underscore the importance of nucleolar structure integrity for organismal fitness.

## Results

### Knocking down class I *rps* genes induces nucleolar reorganization

Ribosomes are composed of two conserved subunits, the large ribosomal subunit (LSU) and the small ribosomal subunit (SSU), which are primarily composed of ribosomal proteins of the large subunit (RPL) and ribosomal proteins of the small subunit (RPS), respectively ^20–22,24,30^. Both RPS and RPL proteins are engaged in various processing and maturation steps of 18S, 5.8S, and 26S rRNAs. Our previous study explored the role of RPL proteins in nucleolar morphology regulation and revealed that the knockdown of class I RPL proteins leads to nucleolar reshaping, in which the spherical nucleolus is transformed into a ring-shaped and vacuole-contained nucleolus ^3^. To investigate whether RPS proteins have similar impacts on nucleolar morphology, we constructed an RRP-8::GFP;mCherry::RPOA-2 strain and treated nematodes with dsRNAs targeting the *rps* genes. RRP-8 is an rRNA processing factor involved in the N1-methyladenosine (m1A) modification of 26S rRNAs ^31^. RPOA-2 is the subunit of RNAP I ^32^. Both RRP-8 and RPOA-2 localize to the nucleolus.

Unlike the distinct separation of FC, DFC, and GC regions observed in mammalian cells, the *C. elegans* nucleolus is organized into two spatially intertwined compartments, the FC and GC regions (Fig. 1A) ^3,12,33^. Upon the knockdown of the 31 *rps* genes, we noticed distinct nucleolar alterations. On the basis of the relative changes in the localization of RRP-8 and RPOA-2, we categorized the *rps* genes into two classes (summarized in Fig. 1C). The distinct spatial separation of RRP-8 and RPOA-2, termed nucleoar demixing. The class I *rps* genes include 13 members, including *rps-1 and rps-4*. RNAi targeting the class I *rps* genes induced the demixing of the RRP-8 and RPOA-2 proteins (Fig. 1B), in which the nucleolus exhibited a three-layer concentric architecture, including the innermost C1 (concentric-spherical nucleolus-1), middle C2 and outermost C3 layers (Fig. 1C). For nomenclature, we named the three-layer nucleolus the concentric-spherical nucleolus. While RRP-8 preferentially localized in the C1 and C3 layers, RPOA-2 exclusively accumulated in the C2 layer after knocking down class I *rps*. The class II *rps* genes include 18 members, including *rps-0 and rps-2*. We did not observe noticeable patterns in the nucleolar structure after RNAi was used to target the class II *rps* genes (Fig. S1A-B).

### Class I RPS and class I RPL proteins act differently in organizing nucleolar architecture

Previously, we showed that upon RNAi targeting class I RPL proteins, RPOA-2-marked nucleoli exhibit a ring-shaped morphology that surrounds the nucleolar vacuole ^3,12^. Here, we also observed a ring-shaped distribution of RPOA-2 after RNAi targeted class I RPS proteins. To distinguish these two rings, we used RNAi to knock down *rps-8* (class I *rps*) and *rpl-14* (class I *rpl*) in the RRP-8::GFP;mCherry::RPOA-2 strain. In control animals not subjected to RNAi, both RRP-8 and RPOA-2 were largely distributed in the spherical nucleolus (Fig. 2A). RNAi targeting *rps-8* induced a three-layer nucleolar architecture, in which RRP-8 accumulated in the innermost C1 and outermost C3 layers, but RPOA-2 accumulated in the middle C2. Knockdown of *rpl-14* resulted in a two-layered nucleolus, in which the nucleolar proteins accumulated in the outside ring and the nucleoplasmic proteins occupied the inside NoV ^3^. Both RRP-8 and RPOA-2 accumulated in the ring-shaped nucleolus but were excluded from the NoV layer (Fig. 2A), suggesting that the two rings induced by *rps-8* and *rpl-14* RNAi are different.

**Figure 2.**
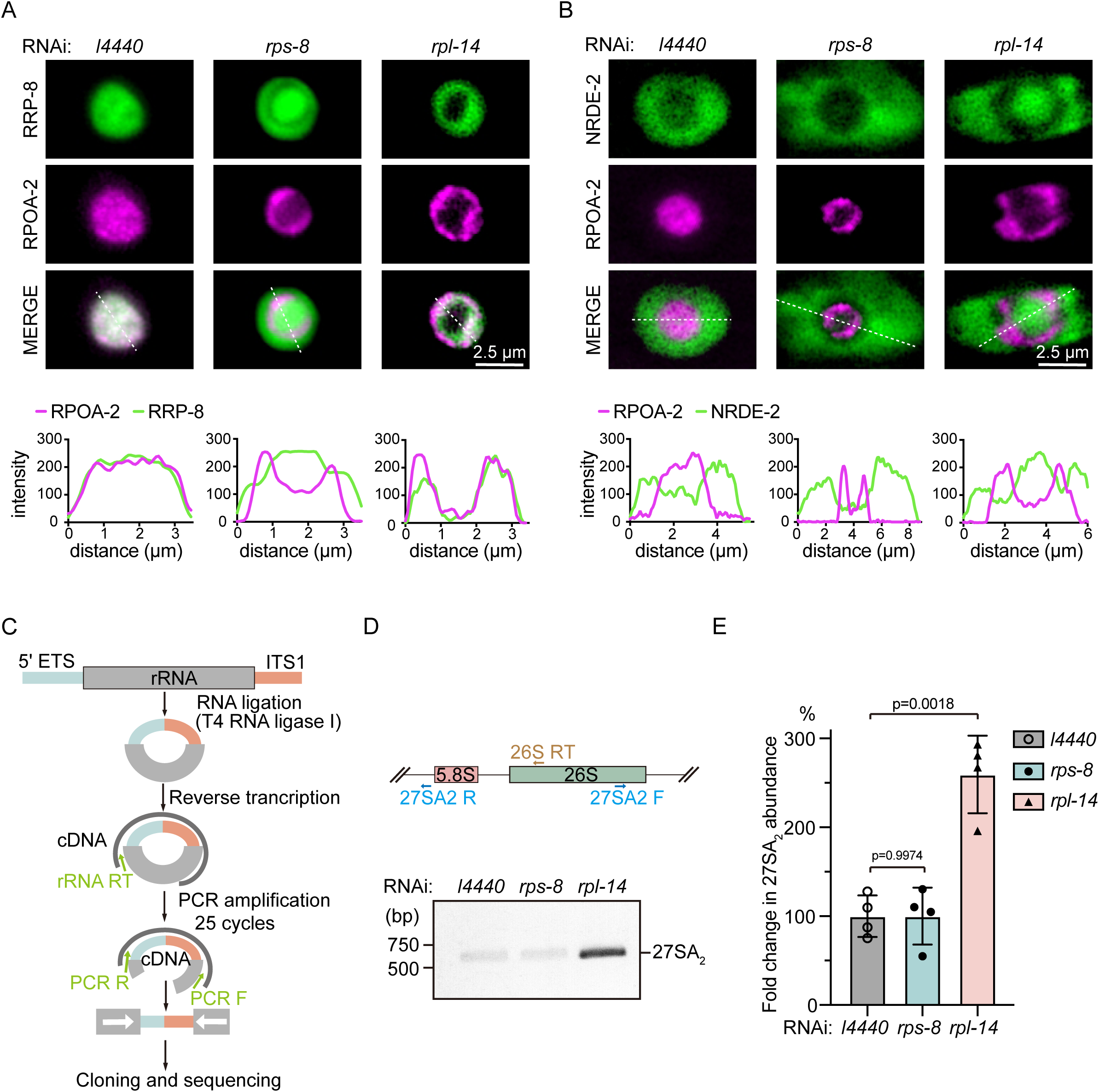
Differential nucleolar structures following the knockdown of class I RPS and class I RPL proteins. (A) (top) Fluorescence microscopy images of *C. elegans* nucleoli in hypodermal cells after the indicated genes were knocked down via RNAi. Scale bar, 2.5 μm. (bottom) Fluorescent density scan of RRP-8::GFP and mCherry::RPOA-2 in the indicated lines via ImageJ. (B) (top) Fluorescence microscopy images of *C. elegans* nucleoli in hypodermal cells after the indicated genes were knocked down via RNAi. Scale bar, 2.5 μm. (bottom) Fluorescent density scan of GFP::NRDE-2 and mCherry::RPOA-2 of the indicated lines via ImageJ. (C) Schematic diagram of the circularized reverse-transcription PCR (cRT-PCR) method. (D) cRT-PCR assay and (E) quantification of 27 SA_2_ pre-rRNA intermediates after knocking down the indicated genes via RNAi. The data are presented as the mean ± SD of four biological replicates. Statistical significance was assessed via a two-tailed t test.

Similarly, we used RNAi to target *rps-8* and *rpl-14* in a GFP::NRDE-2;mCherry::RPOA-2 strain. NRDE-2 is a nucleoplasmic protein that is required for nuclear and nucleolar RNAi ^34–36^. Upon *rps-8* knockdown, NRDE-2 retained its localization in the nucleoplasm. However, NRDE-2 accumulated in both the nucleoplasm and the NoV after *rpl-14* RNAi (Fig. 2B).

To further confirm that the generation of ring-shaped nucleolar structures is different under these two conditions, we used circular reverse transcription PCR (cRT-PCR) ^37^ to assay the 27SA_2_ rRNA intermediate, the increase in which promotes nucleolar reshaping and the formation of NoV ^3^. Knockdown of *rpl-14* induced the accumulation of 27SA_2_ rRNA intermediate, as previously reported ^3^. However, we did not detect the accumulation of 27SA_2_ rRNA after knocking down the class I *rps* gene *rps-8* (Fig. 2C-E). In addition, although vacuole-containing nucleoli are observed in normal growing animals ^3^, concentric-spherical nucleoli are rarely noticeable under normal laboratory culture conditions.

### Distinct demixing behavior of nucleolar proteins during nucleolar structure reorganization

The demixing of RRP-8 and RPOA-2 upon the knockdown of class I RPS proteins suggested that other nucleolar proteins may also exhibit diverse characteristics during the reorganization of the nucleolar architecture. To investigate the subcellular localization of nucleolar proteins during the formation of concentric-spherical nucleoli, we used CRISPR-Cas9 technology to generate single-copy transgenes of a number of nucleolar proteins.

Nucleolar proteins are involved in various steps of rRNA biogenesis, including rRNA transcription, ribosome assembly, pre-rRNA processing and maturation. RPOA-1, RPOA-2, and RPAC-19 are subunits of the RNA polymerase I complex ^32^. DAO-5 is an rRNA transcription factor ^38^. Upon knockdown of *rps-8*, RPOA-1, RPOA-2, RPAC-19, and DAO-5 similarly accumulated in the middle C2 layer (Fig. 1B, 3A, S2A-B). RBD-1, an 18S rRNA processing factor, also localized to the C2 layer upon *rps-8* RNAi (Fig. 3B) ^39^. FIB-1 encodes the *C. elegans* ortholog of human fibrillarin and *Saccharomyces cerevisiae* Nop1p ^40^, which is an essential factor for 18S rRNA processing ^41^. GARR-1 encodes the *C. elegans* ortholog of human GAR1. FIB-1 and GARR-1 normally reside in the FC region ^12^ but accumulated in the middle C2 layer after *rps-8* RNAi (Fig. 3C, S2C). RRP-8 is involved in the N1-methyladenosine (m1A) modification of 26S rRNAs ^31,42^. T06E6.1 is an ortholog of human WDR74 and may be involved in ribosomal large subunit biogenesis ^43^. Both RRP-8 and T06E6.1 accumulated in the C1 and C3 layers upon *rps-8* RNAi (Fig. 1B, 3D). NUCL-1 encodes an evolutionarily conserved protein that is highly homologous to yeast and human nucleolin ^44^. LPD-6 is an ortholog of human PPAN-P2RY11 and yeast Ssf1 and is required for ribosomal large subunit maturation ^45–47^. NUCL-1 and LPD-6 reside in the GC region in control animals and accumulated in the C1 and C3 layers upon *rps-8* RNAi (Fig. 3E, S2D). The results are summarized in Figure 3F and indicate that the factors involved in rRNA transcription, 18S rRNA processing and the fibrillar center accumulate in the C2 layer, whereas the proteins required for 26S rRNA processing, ribosome large subunit assembly and the granular center accumulate in the C1 and C3 layers upon nucleolar demixing.

**Figure 3.**
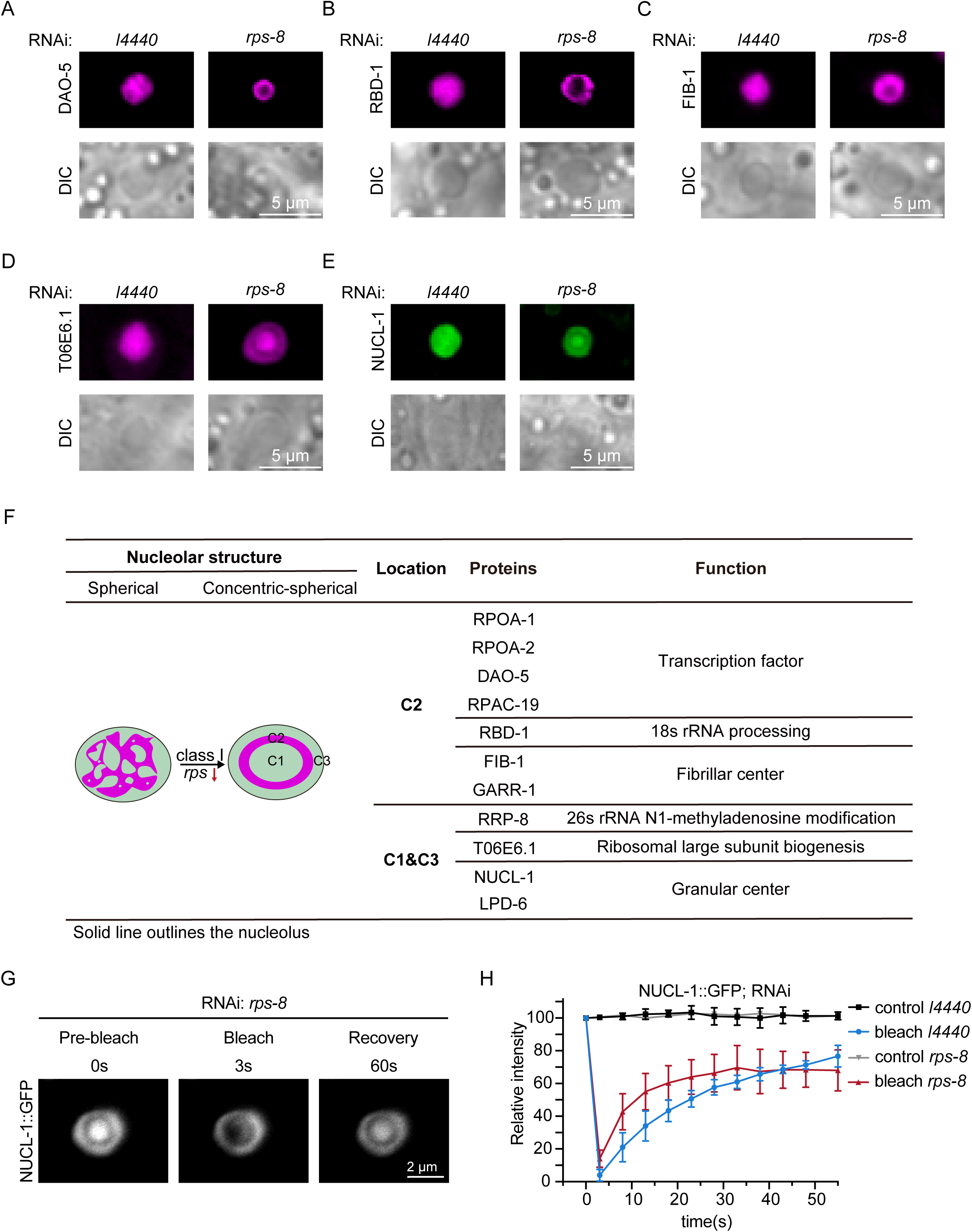
Distinct demixing behavior of nucleolar proteins during nucleolar structure reorganization. (A-E) DIC and fluorescence microscopy images of the indicated transgene after knocking down *rps-8.* Scale bar, 5 μm. (F) Summary of the localization of the indicated proteins after *rps-8* RNAi. (G) FRAP assay of the NUCL-1::GFP transgene in the indicated regions after *rps-8* knockdown. Scale bar, 2 μm. (H) Quantification of FRAP data. Mean ± SD, n = 3 independent animals.

Phase separation of nucleoli is essential for nucleolar function ^7^. We performed a FRAP assay to investigate whether nucleolar demixing alters the mobility of rRNA transcription and processing factors by comparing the mobility of NUCL-1 and GARR-1 in both spherical and concentric-spherical nucleoli. In control and *rps-8*-knockdown animals, both NUCL-1 and GARR-1 exhibited similar mobility (Figs. 3G-H, S2E-F). These data suggested that nucleolar demixing may not affect the mobility of the components for rRNA transcription and processing.

### The Ce.22S rRNA intermediate accumulates upon the knockdown of class I RPS proteins

RPS proteins are 40S ribosome subunits that are involved in 18S rRNA processing and small ribosome assembly ^22,24^. We used cRT-PCR to assay 18S rRNA intermediates via primer sets targeting the external transcribed spacer (ETS) ^48^ upon knocking down *rps* genes (Fig. 4A-B). We identified 4 bands, which were named Ce.20S, Ce.21S, Ce.22S and Ce.23S in reference to the names in yeast (Fig. 4A-B) ^49–51^. The four bands were subsequently cloned and verified via Sanger sequencing (Fig. S3A-D). There are two putative TATA-like sequences: one located 420 bp upstream and the other 396 bp upstream of the 18S rRNA-coding sequence ^39^. We detected that transcription likely starts downstream of the second TATA-like sequence, which we designate the -394(18S) site (Fig. S3A-B). In addition, we mapped cleavage sites III and IV to positions +245(18S) and +376(18S), respectively (Fig. S3C-D). These regions within ITS1 correspond to those previously defined as “III” and “IV” by *Saijou* and *Ushida* et al. via northern blotting and primer extension analyses, respectively ^39,52^.

**Figure 4.**
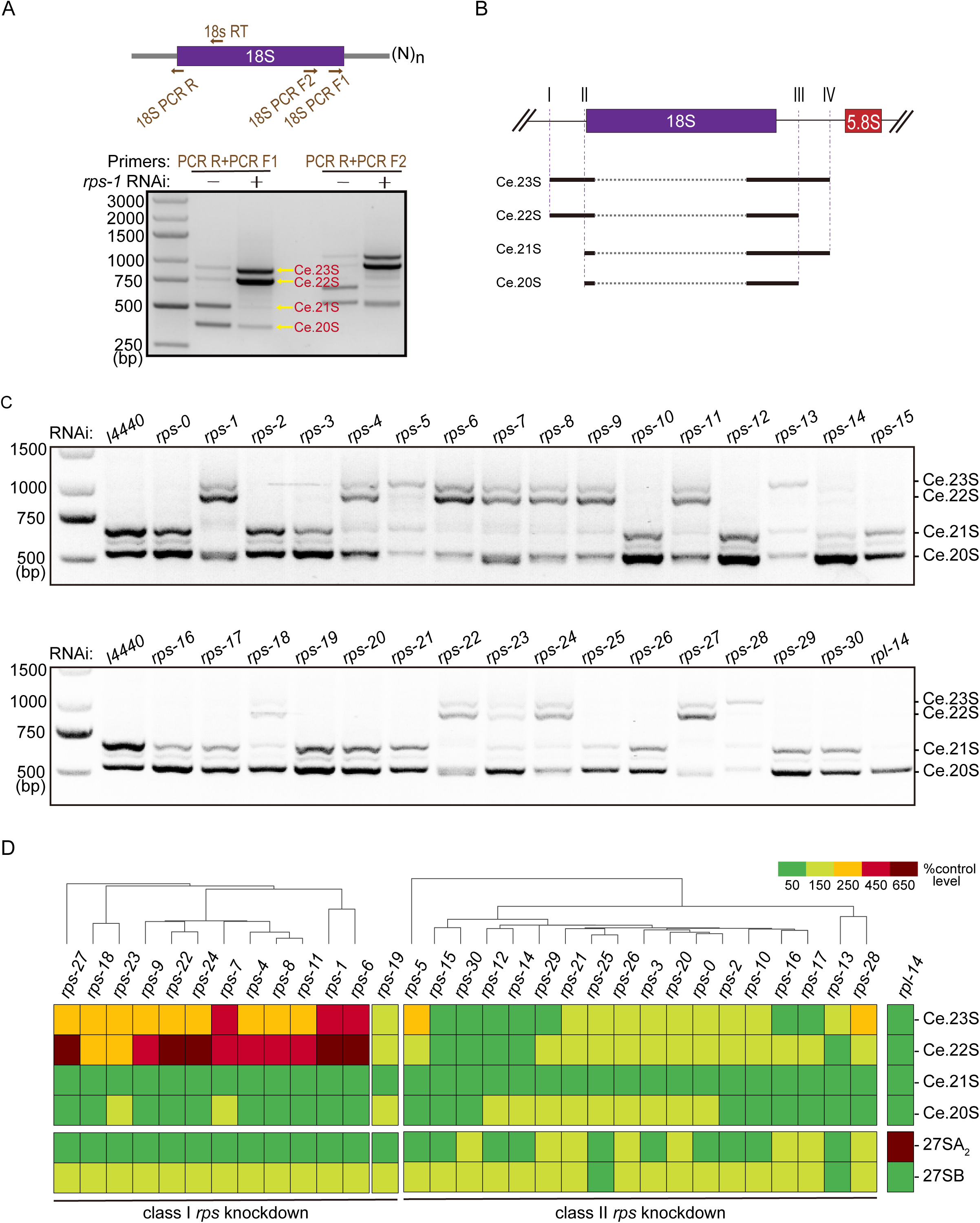
Knocking down class I RPS proteins induces the accumulation of the Ce.22S rRNA intermediate. (A) cRT-PCR assay and (B) schematic diagram of 18S pre-rRNA intermediates. (C) cRT-PCR assay of 18S pre-rRNA intermediates after knocking down the indicated *rps* genes via RNAi. (D) Quantification of 18S and 26S pre-rRNA intermediates (Fig. 4C and S3E, respectively) via ImageJ after knocking down the indicated *rps* genes via RNAi and normalization to the corresponding intermediates in *L4440* RNAi controls. Hierarchical clustering analysis was performed on the basis of the normalized values.

Strikingly, the knockdown of class I *rps* consistently led to the accumulation of both Ce.22S and Ce.23S pre-rRNAs, whereas the knockdown of class II *rps* had a subtle effect on their levels (Fig. 4C-D, S3E). A few other *rps* genes, including *rps-5, rps-13,* and *rps-28,* which are not class I *rps genes*, still accumulated Ce.23S pre-rRNAs but not Ce.22S pre-rRNAs. Taken together, these data implied that inappropriate accumulation or processing of Ce.22S rRNA may be the trigger for the formation of concentric-spherical nucleoli. An exception is *rps-19*, the knockdown of which induced a demixing phenomenon without the accumulation of Ce.22S rRNA intermediate (Fig. S1B, 4C). The reason is still unknown.

Interestingly, the classification of *rps* genes in *C. elegans* is coincidentally consistent with the classification of *rps (i-rps/p-rps)* genes in human cells ^53^. In HeLa cells, the knockdown of i-RPS results in the accumulation of 45S and 30S pre-rRNAs because of a failure of the processing steps within the 5′ external transcribed spacer (5′ ETS) and internal transcribed spacer 1 (ITS1). In *C. elegans*, the accumulation of Ce.22S rRNA also arises from defective cleavage of the 5′ ETS. These findings imply the critical roles of RPS proteins in rRNA biogenesis, ribosome assembly and nucleolar architecture.

Taken together, these data suggest that the rRNA processing steps or intermediates are involved in the maintenance of nucleolar architecture.

### Inhibition of rRNA transcription induces the formation of sandwich-like nucleoli

We then tested whether and how the reduction in rRNA levels impacts the nucleolar architecture by using actinomycin D to inhibit rRNA transcription. Our previous work revealed that actinomycin D inhibits the accumulation of 27SA_2_ rRNA intermediates and prevents the formation of NoV ^3^.

We treated the RRP-8::GFP;RPOA-2::mCherry strain with 30 µg/ml actinomycin D and visualized the nucleolar substructure. Upon actinomycin D treatment, the pre-rRNA levels decreased (Fig. S4A). RRP-8 and RPOA-2 demixed and resembled a sandwich-like structure (Fig. 5A, S4B), which is distinct from the concentric-spherical nucleolus, yet maintained a similar layered organization, with RRP-8 occupying the outside layers S1&S3 (sandwich-like nucleoli-1& -3) and RPOA-2 accumulating in the middle S2 layer. The actinomycin D-induced nucleolar protein demixing is similar to the nucleolar architecture reorganization triggered by class I *rps* knockdown (Fig. 5B-G, S4C-F and summarized in Fig. 5B). The factors involved in rRNA transcription, 18S rRNA processing and FC organization, such as GARR-1, FIB-1, RPOA-1, RPOA-2, DAO-5, RPAC-19, and RBD-1, are localized to the sandwich core (S2). The outer layers of the sandwich-like nucleolar structure (S1 & S3) were enriched with proteins required for 26S rRNA processing, large subunit assembly, and GC organization, including NUCL-1, LPD-6, RRP-8, and T06E6.1. Therefore, we speculate that active rRNA transcription and the presence of particular rRNA intermediates are essential for the maintenance of nucleolar architecture.

**Figure 5.**
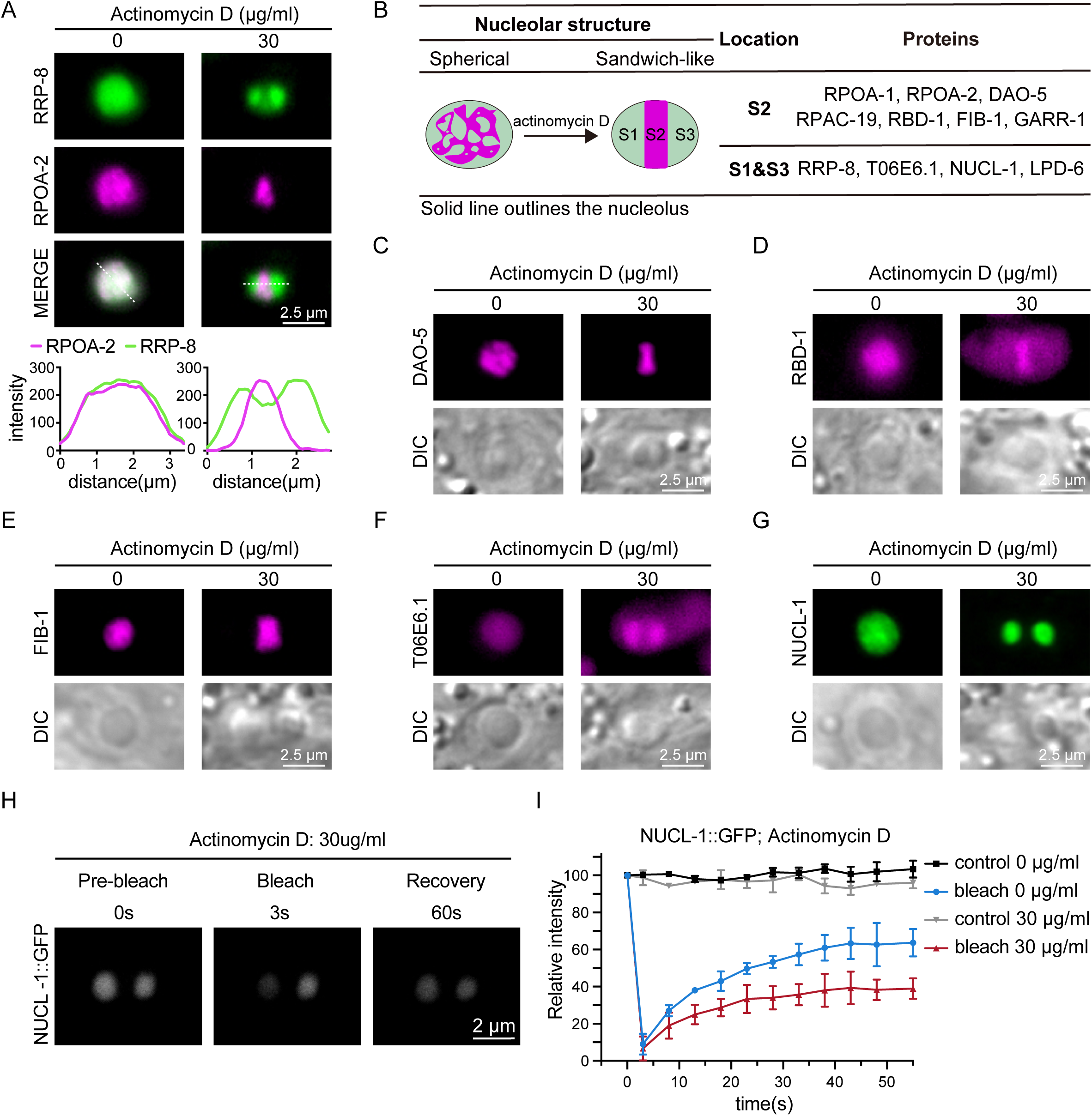
Inhibition of rRNA transcription induces the formation of sandwich-like nucleoli. (A) (top) Fluorescence microscopy images of *C. elegans* nucleoli in hypodermal cells after actinomycin D treatment for 48 h. Scale bar, 2.5 μm. (bottom) Fluorescence density scan of RRP-8::GFP and mCherry::RPOA-2 in the indicated lines via ImageJ. (B) Summary of the localization of the indicated proteins after actinomycin D treatment for 48 h. (C-G) DIC and fluorescence microscopy images of the indicated transgenes after actinomycin D treatment for 48 h. Scale bar, 2.5 μm. (H) FRAP assay of the NUCL-1::GFP transgene in the indicated regions after 30 µg/ml actinomycin D treatment for 48 h. Scale bar, 2 μm. (I) Quantification of FRAP data. Mean ± SD, n = 3 independent animals.

We examined protein mobility via the FRAP assay and observed a slight decrease in the mobility of NUCL-1 and GARR-1 upon actinomycin D treatment (Fig. 5H-I, S4G-H), suggesting a slightly more compact structure of the nucleolus upon the inhibition of rRNA transcription.

### Spatial distribution of rDNA during structural reorganization of nucleoli

In addition to proteins and rRNAs, rDNA has been shown to play critical roles in nucleolar architecture as the nucleolar organizer region (NOR) ^2^. We then investigated the subcellular localization of rDNA during nucleolar structural alterations. To achieve live-cell labeling of the rDNA genome, we employed the lacI-lacO system ^54–58^ and ectopically inserted the lacI::tagRFP transgene into the *C. elegans* genome. Moreover, the lacO sequence was integrated into the rDNA genome via CRISPR-Cas9 technology to enable the visualization of the rDNA array by lacI::tagRFP (Fig. 6A). Notably, the *C. elegans* genome encodes approximately 55 copies of a 7.2 kb tandem repeat of rDNA on LG I to express pre-rRNAs ^15^. Owing to the length limitation of DNA sequencing, we are unable to determine in which specific rDNA unit the lacO sequence was inserted. To investigate the positioning of rDNA, we used a zoning assay ^58–60^ and examined the position of rDNA foci relative to zones C1-C3 and S1-S3 under a fluorescence microscope.

**Figure 6.**
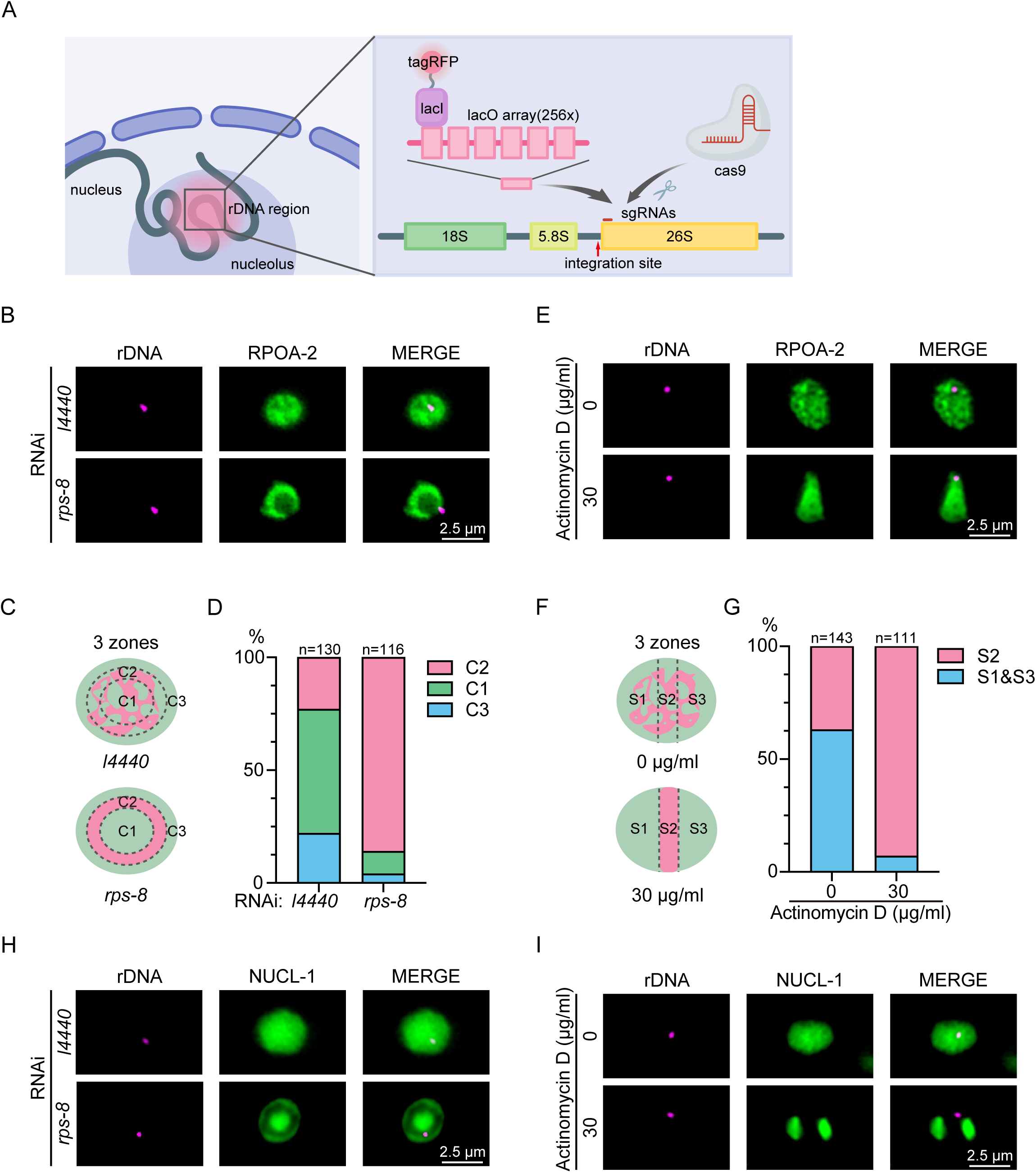
Localization of rDNA upon nucleolar structure reorganization. (A) Schematic diagram of the lacI::tagRFP-lacO system used for rDNA labeling. lacO assay: 17 bp lacO + 19 bp spacer. (B) Fluorescence images of rDNA (labeled with lacI::tagRFP) and GFP::RPOA-2 after *rps-8* knockdown. Scale bar, 2.5 μm. RPOA-2 marks the C2 region. (C) Zoning assay for rDNA reporter distribution on the basis of the C1-C3 regions as defined in Figure 1C. (D) Quantification of the subnucleolar distribution of rDNA. *n,* number of nucleoli. (E) Fluorescence microscopy images of rDNA (labeled with lacI::tagRFP) and GFP::RPOA-2 after actinomycin D treatment. Scale bar, 2.5 μm. RPOA-2 marks the S2 region. *n*, number of nucleoli. (F) Zoning assay using the S1-S3 regions from Figure 5C. (G) Quantification of the subnucleolar distribution of rDNA. *n*, number of nucleoli. (H) Fluorescence microscopy images of rDNA (labeled with lacI::tagRFP) and NUCL-1 after *rps-8* knockdown. Scale bar, 2.5 μm. NUCL-1 marks the C1&C3 region. (I) Fluorescence microscopy images of rDNA (labeled with lacI::tagRFP) and NUCL-1 after actinomycin D treatment. Scale bar, 2.5 μm. NUCL-1 marks the S1 & S3 region.

Upon *rps-8* knockdown, rDNA foci significantly accumulated in the C2 zone, in which RNAP I transcription, 18S processing, and FC region proteins were preserved (Fig. 6B-D). In actinomycin D-induced sandwich-like nucleoli, approximately 90% of rDNA foci were positioned within the S2 zone (Fig. 6E-G), in which the RNAP I transcription, 18S processing, and FC region proteins were enriched as well. We did not observe pronounced accumulation of rDNA in the C1 & C3 or S1 & S3 regions (Fig. 6H-I).

Therefore, rDNA remains colocalized with rRNA transcription and 18S rRNA processing factors during nucleolar structural reorganization.

### NUCL-1 is required for nucleolar demixing and development

NUCL-1 in *C. elegans* encodes an evolutionarily conserved protein with high homology to nucleolin in yeast and humans. It localizes to the GC region of the nucleolus, and its N-terminal domain contains a long intrinsically disordered region (IDR) with a GAR/RGG motif that is essential for its subnucleolar compartmentalization ^3,4,12^. Our previous work revealed that NUCL-1 is required for nucleolar reshaping and NoV formation ^3^. Here, we tested whether NUCL-1 is also required for nucleolar demixing and the formation of concentric-spherical nucleoli.

The deletion of *nucl-1* significantly suppressed the formation of *rps-8* knockdown-induced concentric-spherical nucleoli (Fig. 7A-B). Moreover, the RNAi knockdown efficiency of *rps-8* was not compromised in the *nucl-1* mutant, suggesting that RNAi activity was not compromised in the *nucl-1* mutant (Fig. S5A). The FRAP assay revealed that the mobility of the spherical nucleoli in *nucl-1;rps-8* animals was similar to that in control animals (Fig. S5B). cRT-PCR revealed that the deletion of NUCL-1 blocked the *rps-8*(RNAi)-induced accumulation of Ce.22S rRNA (Fig. 7C‒D), further supporting the correlation between the accumulation of Ce.22S rRNA and nucleolar demixing.

**Figure 7.**
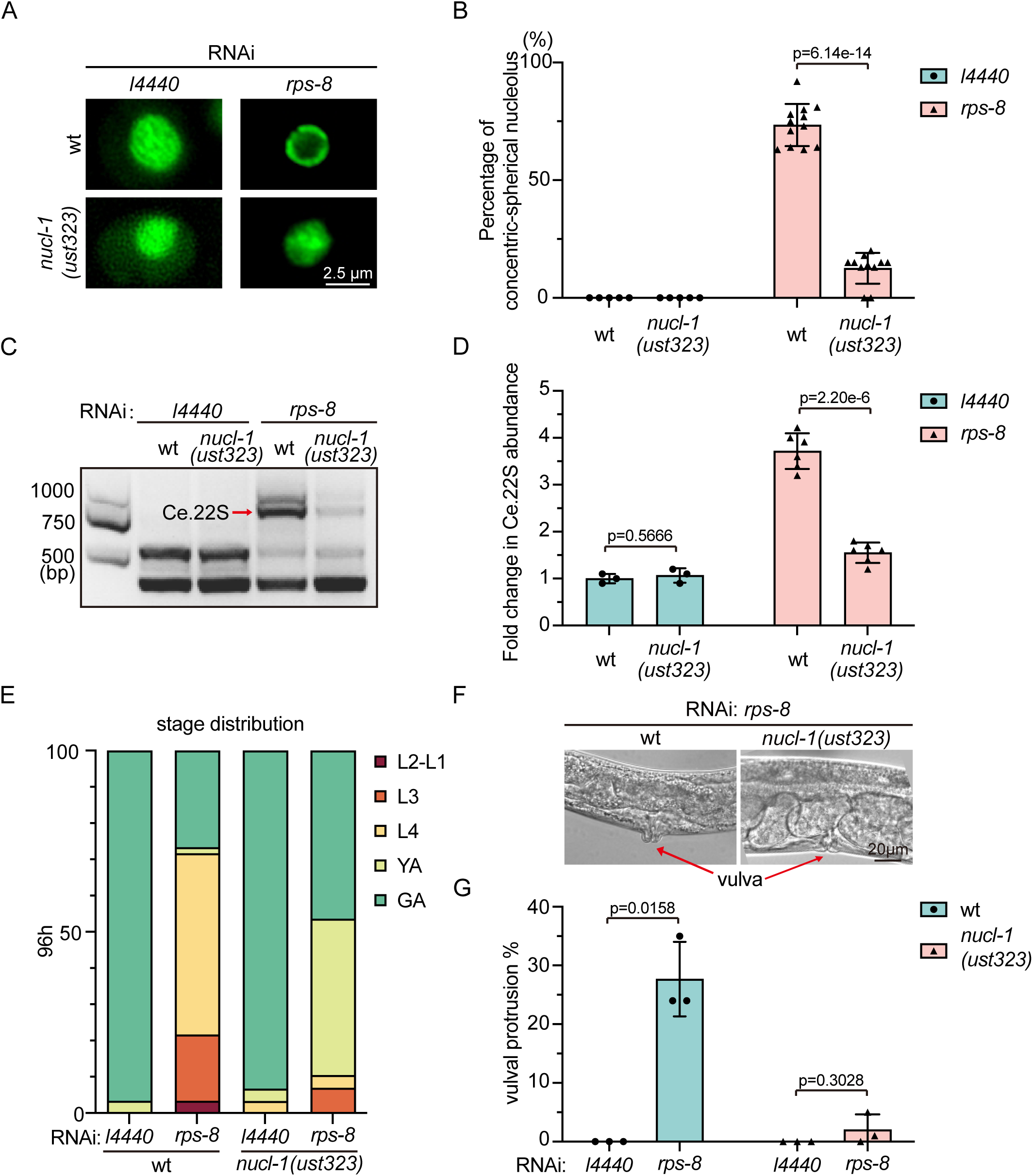
NUCL-1 is required for nucleolar structure reorganization. (A) Fluorescence microscopy images of *C. elegans* nucleoli in hypodermal cells after the indicated genes were knocked down via RNAi in wild-type (wt) and *nucl-1* mutants. Scale bar, 2.5 μm. (B) Quantification of concentric-spherical nucleoli cells in which the indicated genes were knocked down via RNAi in wt and *nucl-1* mutants. Mean ± SD, n ≥5 independent animals. A two-tailed t test was performed to determine statistical significance. (C) cRT-PCR assay and (D) quantification of the Ce.22S rRNA intermediate in *nucl-1* mutants upon knockdown of the indicated genes via RNAi. (C) The red arrow indicates the Ce.22S rRNA intermediate. (D) Data are presented as the mean ± SD of three or six biological replicates. A two-tailed t test was performed to determine statistical significance. (E) Developmental stage assessment of the indicated animals after 96 h at 20°C; n ≥18 animals. (F) DIC images and (G) quantification of animals whose vulva protruded upon knocking down *rps-8* in wt and *nucl-1* mutants. (F) The red arrow indicates the vulva. Scale bar, 20 μm. (G) Mean ± SD, n ≥18 animals. A two-tailed t test was performed to determine statistical significance.

To investigate the biological roles of NUCL-1, we examined the developmental progress of *C. elegans* upon *rps-8* knockdown with or without *nucl-1*. The depletion of *rps-8* delayed the delelopment of nematodes, which could be partially rescued by *nucl-1* mutation (Fig. 7E). *rps-8* knockdown induced the protruding vulvar phenotype, which could also be alleviated by additional *nucl-1* mutation (Fig. 7F-G). Notably, the deletion of *nucl-1* also prevented the formation of actinomycin D-induced sandwich-like nucleolar structures (Fig. S6A-B), but *nucl-1* mutation failed to restore pre-rRNA levels (Fig. S6C) or alter nucleolar fluidity (Fig. S6D), suggesting that NUCL-1 may play diverse roles in regulating nucleolar structure and function in response to different growth conditions.

Together, these results suggest that NUCL-1 may regulate nucleolar structure reorganization and facilitate development in *C. elegans* when rRNA production is impaired.

## Discussion

The nucleolus is a dynamic, multifunctional organelle that serves as the central hub for rRNA synthesis and ribosome assembly. It also participates in various cellular stress responses, including DNA damage, heat shock, transcriptional inhibition, and defects in ribosome biogenesis. Perturbations in rRNA biogenesis and ribosome production often induce marked changes in nucleolar morphology and composition ^61,62^. We previously showed that the knockdown of class I *rpl* genes in *C. elegans* leads to abnormal accumulation of 27SA_2_ rRNA intermediate, reshapes nucleolar morphology, and results in the formation of ring-shaped nucleoli and nucleolar vacuoles (NoVs) ^3^. Here, we further investigated whether and how other perturbations of nucleolar processes influence nucleolar architecture, including pre-18S rRNA processing and rRNA transcription. We found that the knockdown of a specific subset of RPS proteins led to the accumulation of the Ce.22S rRNA intermediate, resulting in the formation of concentric-spherical nucleoli.Similarly, the inhibition of rRNA transcription leads to the formation of another three-layered architecture, a sandwich-like nucleoli. During both modes of nucleolar structure reorganization, nucleolar proteins undergo demixing and spatially redistribute into distinct subnucleolar regions. Furthermore, we found that *nucl-1* knockout prevents nucleolar demixing and nucleolar structure reorganization. Collectively, these results reveal that diverse impairments in rRNA processing can elicit distinct alterations in nucleolar architecture, underscoring the essential contribution of rRNA biogenesis to the structural homeostasis of the nucleolus (Fig. S7).

### RPS-mediated rRNA processing is evolutionarily conserved

rRNA constitutes approximately 80% of the total RNA within a cell. The production of rRNA is a complex, multistage process that involves the modification, cleavage and folding of rRNAs with the assistance of various assembly factors, including ribosomal proteins ^48,63^. In this study, via a circular PCR-based method, we mapped the precise cleavage sites of pre-18S rRNA and identified a series of intermediates. These cleavage sites are indeed located in reported *C. elegans* rRNA processing regions, as revealed by northern blotting and primer extension methods ^39,52^. We further systemically examined the functions of RPS proteins in pre-18S rRNA processing and revealed that the depletion of a specific subset of RPS proteins led to failure of cleavage in both the 5*’* ETS and ITS1 regions, resulting in the accumulation of Ce.23S and Ce.22S rRNA intermediates. Interestingly, in mammalian cells, the depletion of the homologs of these RPS proteins, which are classified as i-RPSs (short for initiation-RPSs), also leads to defects in the cleavage of the 5*’* ETS and ITS1 regions, resulting in the accumulation of 30S rRNA intermediate ^53,64^. Although both the cleavage sites of rRNA and the sequences of pre-rRNA intermediates in *C. elegans* are not exactly the same as those in mammalian cells are, these findings suggest that the functions of RPS proteins in pre-rRNA processing are highly conserved across species.

### rRNA processing intermediates serve as dictators of nucleolar architecture

As a multilayered membraneless structure within the cell, the nucleolus is primarily responsible for rRNA production, processing and ribosomal assembly. Current models posit that the multilayered structure of the nucleolus may facilitate the transcription and successive processing of rRNAs ^7,65^. Nevertheless, whether and how rRNA transcription and processing regulate the layered organization of the nucleolus remain poorly understood. Recent studies have revealed that rRNA may undergo directed movement rather than passive diffusion in the nucleolus and that the sequential maturation of rRNA promotes its outward flux, suggesting that the sequential processing and advective flow of rRNA may underlie the nucleolar form ^18,66^. Moreover, in our previous study, we showed that defects in 27SA_2_ rRNA processing lead to the formation of vacuole-containing nucleoli in *C. elegans* ^3^, implying that proper rRNA processing may help shape the spatial structure of the nucleolus. Here, by systemically examining the functions of RPS proteins in the organization of nucleolar structure and pre-18S rRNA processing, we further investigated how pre-rRNA processing contributes to the multilayered organization of the nucleolus in *C. elegans*. We revealed that the depletion of class I RPS proteins leads to the formation of a concentric three-layered nucleolar architecture and the accumulation of Ce.23S and Ce.22S rRNAs.

Interestingly, Ce.23S and Ce.22S rRNAs have the same 5*’* ETS sequence, but only differ in the 3*’* ITS1 sequence. However, enriching only Ce. 23S rRNA does not induce nucleolar structural reorganization, suggesting that Ce.22S rRNA with a shorter 3*’* ITS1 sequence may play a more critical role in the reorganization of the nucleolar structure. A recent study in human cells revealed that defects in SSU processing via the knockdown of U3 snoRNAs alter the ordering of the nucleolar structure, resulting in inside-out nucleoli, in which the GC region relocates to the nucleolar center ^66^. Although the multilayered structure of the nucleolus in nematodes is different from that in human cells, these findings suggest a potential link between specific rRNA processing events and the structural organization of the nucleolus and indicate that the regulation of nucleolar structure by pre-18S rRNA intermediates may be conserved across species. Further investigation of the effects of the disruption of specific rRNA processing steps on nucleolar structure in different organisms, including human cells, may help reveal how early pre-rRNA processing helps organize nucleolar architecture.

rRNAs are essential for sustaining nucleolar fluidity. In both *C. elegans* and mouse embryos, the inhibition of transcription leads to reduced rRNA output and a phase transition in the GC region, marked by slower diffusion and increased viscosity ^67^. Changes in the phase separation properties of nucleolar subcompartments have been shown to alter nucleolar morphology and function ^68^. We propose that the presence of rRNA processing intermediates may dictate the local microenvironment of the nucleolus or interact with certain nucleolar proteins in a way that changes the organization of the nucleolar structure. In human cells, the inhibition of rRNA transcription leads to the formation of nucleolar caps, reflecting segregation or demixing of nucleolar components. In *C. elegans*, transcriptional inhibition results in the formation of sandwich-like nucleoli. This similarity between nematodes and mammals suggests that the underlying mechanism might be conserved: reduced rRNA levels alter the properties of nucleolar proteins, triggering nucleolar architecture reorganization. Therefore, rRNA is not merely a passive output of nucleolar function but also an active organizer of the nucleolar structure.

Taken together, our work revealed that changes in nucleolar morphology are tightly associated with the status of rRNA transcription and processing. However, it remains unclear whether the changes in nucleolar architecture are a manifestation of the inappropriate accumulation of rRNA processing intermediates or a result of altered viscoelastic properties of unprocessed rRNA. Given rRNA’s pivotal role in shaping nucleolar architecture, developing a new live rRNA labeling system is essential to better reveal its spatial organization and function within the nucleolus. Moreover, in vitro nucleolar reconstruction experiments subsidized with distinct synthesized rRNA intermediates will further help in understanding nucleolar structure organization, including but not limited to reshaping, mixing and demixing behaviors.

### NUCL-1 is required for nucleolar structure reorganization

NUCL-1 is a homolog of human nucleolin (NCL) and the longest RGG domain-containing protein in the nucleolus ^12^. Previous studies have shown that *nucl-1* knockout leads to delayed development after 5 generations ^12^, yet its function remains largely unclear. Here, we showed that NUCL-1 plays a crucial role in promoting nucleolar reorganization in response to impaired rRNA transcription and processing. Furthermore, we observed that *nucl-1* knockout partially reversed the *rps-8* depletion-induced delay in growth. Moreover, *nucl-1* mutation prevented the accumulation of the Ce.22S rRNA intermediate induced by *rps-8* knockdown. We speculated that the nucleolar architecture is not only shaped by rRNA processing but also feeds back to influence rRNA maturation via NUCL-1. Notably, NUCL-1 may act as a checkpoint of rRNA processing, thereby ensuring the fidelity of ribosome biogenesis.

Intriguingly, haploinsufficiency of ribosomal protein (RP) genes leads to ribosomopathies in humans, such as Diamond-Blackfan anemia (DBA) ^69^ and acute myeloid leukemia (AML), for which effective therapies are still lacking. These ribosomopathies are frequently linked to dysregulated rRNA processing ^70–74^. For example, mutation of the DBA gene *rps7* leads to the accumulation of the 30S rRNA intermediate ^73^, indicating failure of 5′ ETS cleavage. Our work highlights NUCL-1/NCL as a potential regulatory node in this process and underscores the importance of nucleolar architecture in maintaining rRNA and ribosome homeostasis. Further investigation into the mechanistic interplay between rRNA processing, nucleolar morphology, and developmental processes may yield novel insights and therapeutic avenues for ribosomopathies.

## Materials and Methods

### Strains

The Bristol strain N2 was used as the standard wild-type strain. All strains were grown at 20°C unless otherwise specified. The strains used in this study are listed in Table S1.

### RNAi

RNAi experiments were performed at 20°C by placing synchronized embryos on feeding RNAi plates as previously described ^75^. HT115 bacteria expressing the empty vector L4440 (a gift from A. Fire) were used as controls. Bacterial clones expressing double-stranded RNAs (dsRNAs) were obtained from the Ahringer RNAi library ^76^ and sequenced to verify their identity. Some bacterial clones, which are listed in Table S2, were constructed in this work.

### Imaging

Images were acquired via a Leica DM4B microscope with a Leica Thunder image-processing system. All worms were imaged at the L3-L4 stage unless otherwise specified.

### Construction of plasmids and transgenic strains

For the ectopic transgenes, the promoter and CDS regions and UTRs were amplified from N2 genomic DNA. A GFP::3xFLAG sequence was PCR amplified from SHG326 genomic DNA. The mCherry coding sequence was amplified from PFCJ90. The vector fragment was PCR amplified from the plasmid pSG274. These fragments were joined together by Gibson assembly to form the repair plasmid with the ClonExpress MultiS One Step Cloning Kit (Vazyme Biotech, Nanjing, China, Cat. No. C113-01/02). The transgene was integrated into *C. elegans* chromosomes I and II via a modified counterselection (cs)-CRISPR method ^77,78^. The sequences of the primers are listed in Table S3.

For in situ knock-in transgenes, the 3xFLAG::GFP coding region was PCR amplified from shg1248 genomic DNA. The GFP:3xFLAG coding region was PCR amplified from shg2123 genomic DNA. The mCherry coding region was PCR amplified from shg1660 genomic DNA. The tagRFP coding region was PCR amplified from YY1446 genomic DNA. The homologous left and right arms (1.5 kb) were PCR amplified from N2 genomic DNA. The backbone was PCR amplified from the plasmid pCFJ151. All these fragments were joined together by Gibson assembly to form the repair plasmid with the ClonExpress MultiS One Step Cloning Kit (Vazyme Biotech, Nanjing, China, Cat. No. C113-01/02). This plasmid was coinjected into N2 animals with three sgRNA expression vectors, 5 ng/mL pCFJ90 and the Cas9 II-expressing plasmid. The sequences of the primers are listed in Table S4.

### Actinomycin D treatment

Actinomycin D (MedChemExpress no. HY-17559, CAS:50-76-0) was prepared at 20 mg/mL in DMSO as a stock solution. Each 10.5 μL of actinomycin D stock solution was diluted with 300 μL of concentrated OP50 and layered onto NGM plates. Synchronized embryos were placed onto the seeded plates and grown for 48 h before imaging.

### Fluorescence recovery after photobleaching (FRAP)

FRAP experiments were performed via a Zeiss LSM980 laser scanning confocal microscope at 20°C. Worms were anesthetized with 2 mM levamisole. A region of interest was bleached with 100% laser power for 3-4 s, and the fluorescence intensities in these regions were collected every 5 s and normalized to the initial intensity before bleaching. For analysis, image intensity was measured as the mean and further analyzed with GraphPad Prism 10.0 software.

### cRT-PCR

L3 stage worms were incubated with TRIzol (Invitrogen) reagent followed by seven quick liquid nitrogen freeze‒thaw cycles. RNA was precipitated with isopropanol followed by DNaseI digestion (Thermo Fisher). Two micrograms of total RNA was circularized with a T4 RNA Ligase 1 Kit (M0204. NEB) and then purified with TRIzol reagent followed by isopropanol precipitation. The circularized RNA was reverse transcribed via the GoScript Reverse Transcription System (Promega #A5001). PCR was performed via 32 cycles of 2X Phanta Flash Master Mix (Vazyme, P520-AA). The primer sets used in this work are listed in Table S5.

### Quantitative real-time PCR

All quantitative real-time PCR (qPCR) experiments were performed via a Roche system. cDNA was quantified with SYBR Green Master Mix (Vazyme, Q111-02), and qPCR was performed according to the vendor’s instructions. The primer sets used in this work are listed in Table S6.

### Stage distribution

Five gravid adults were allowed to lay eggs for 1 h on the indicated RNAi plates and then removed. The developmental progress was then monitored after 96 h at 20°C.

### Statistics

Statistics Boxplots are presented with medians and minimums and maxima. Bar graphs with error bars are presented, which present the means and standard deviations (SDs). All experiments were conducted with independent *C. elegans* animals at the indicated number of times (N). Statistical analysis was performed with a two-tailed Student’s t test.

## Acknowledgments

We are grateful to the members of the Guang laboratory for their comments and suggestions. We are grateful to the International *C. elegans* Gene Knockout Consortium and the National Bioresource Project for providing the strains. Some strains were provided by the CGC, which is funded by the NIH Office of Research Infrastructure Programs (P40 OD010440).

## Author contributions

S.G., X.F., X.C., and J.C. conceptualized the research; Xinya Huang, X.C., C.Z., X.F., and S.G. designed the research; J.C., L.L., D.X., Y.K., A.Z., M.H., Xiaona Huang, X.Z., W.N. and Xinya Huang performed the research; J.C., L.L., D.X., and Y.K. contributed new reagents; J.C., X.C., X.F., and S.G. wrote the paper.

---

**Figure S1.**
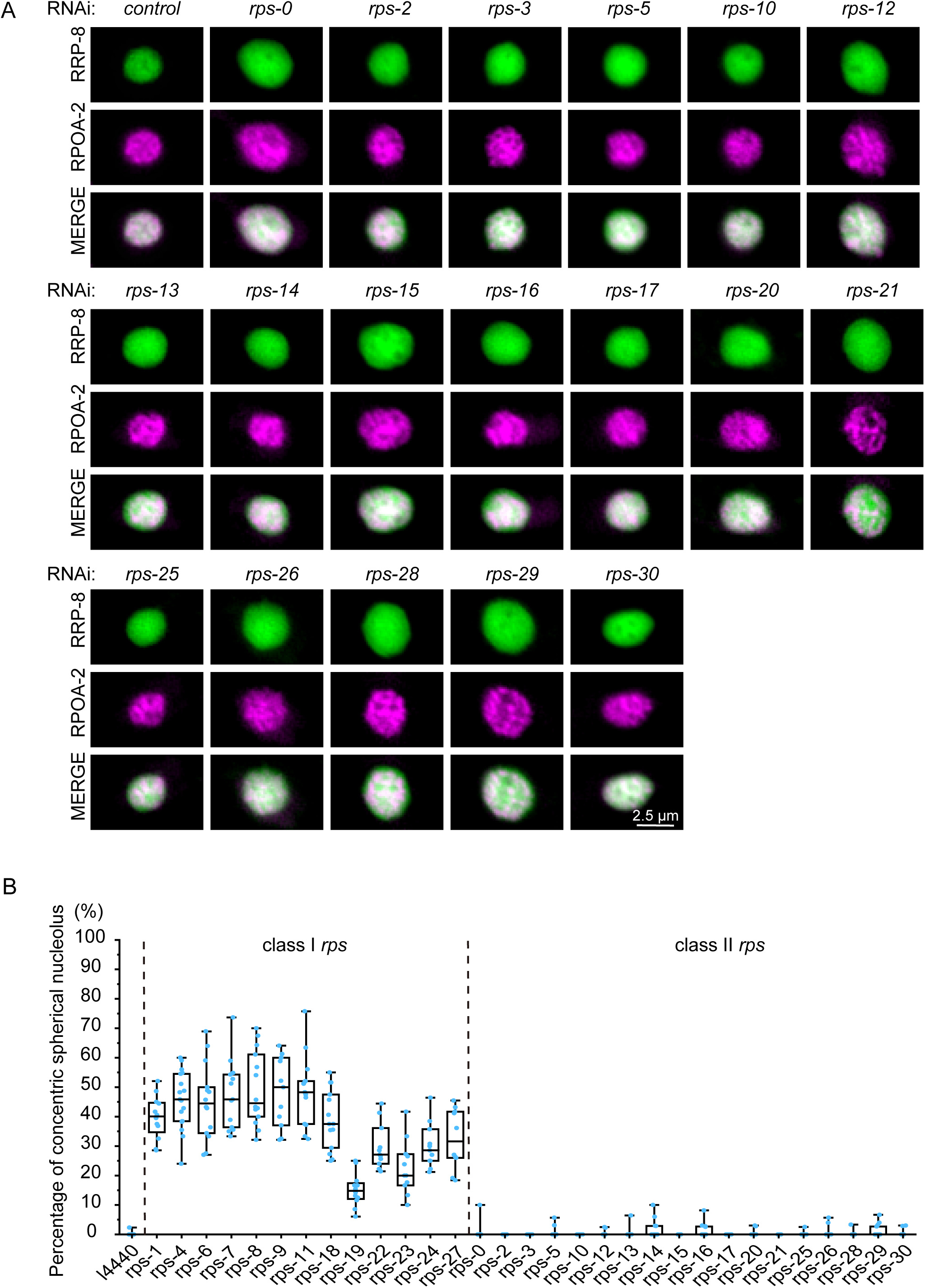
Knocking down class I *rps* genes induces nucleolar reorganization. (A) Fluorescence microscopy images of *C. elegans* nucleoli in hypodermal cells after the indicated genes were knocked down via RNAi. Scale bar, 2.5 μm. (B) Quantification of concentric-spherical nucleoli after RNAi was used to knock down the indicated genes. The percentage of animals in which **at least three hypodermal cells** presented concentric-spherical nucleoli was quantified. n ≥ 11 animals.

**Figure S2.**
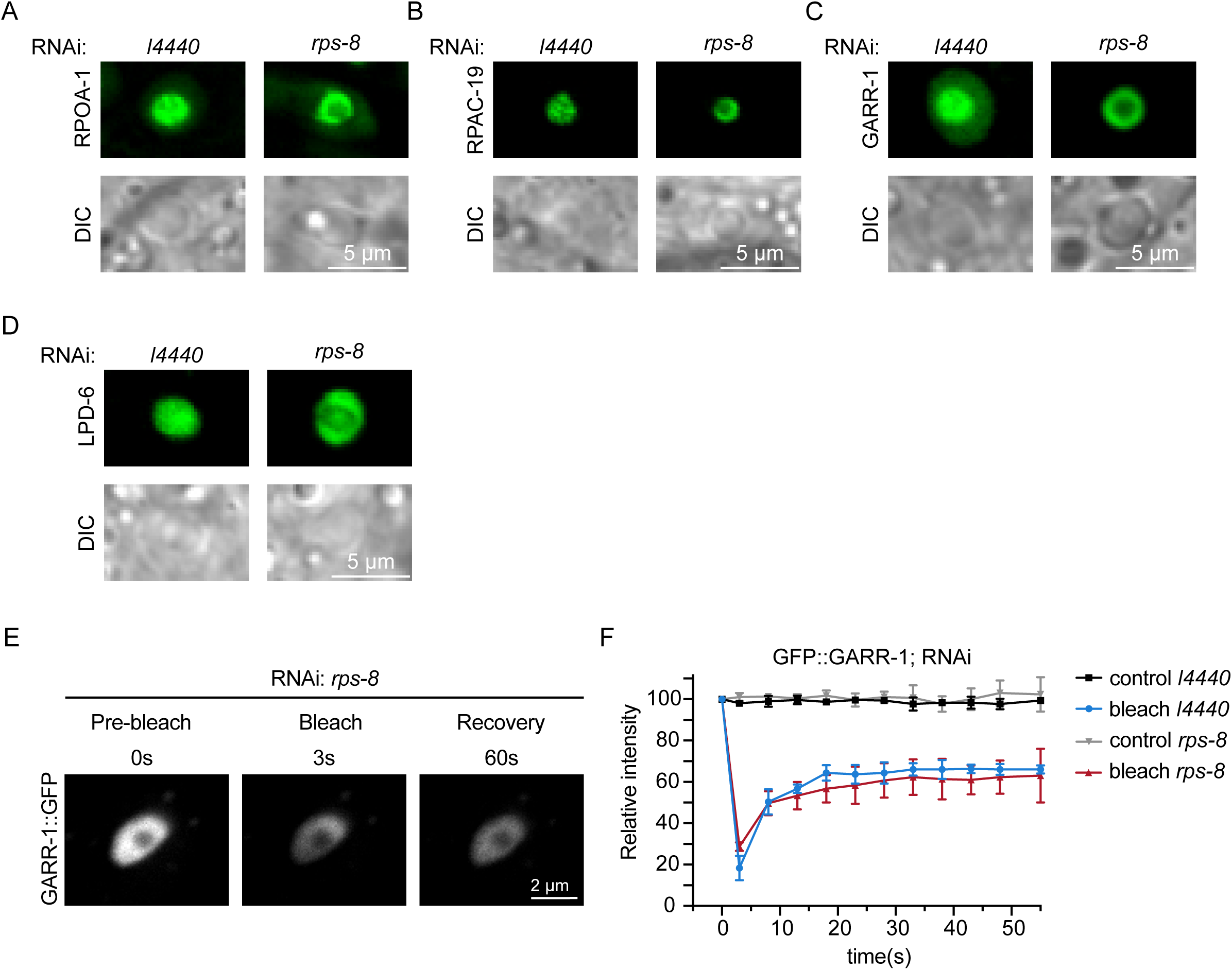
Distinct demixing behavior of nucleolar proteins during nucleolar structure reorganization. (A-D) DIC and fluorescence microscopy images of the indicated transgene after knocking down *rps-8.* Scale bar, 5 μm. (E) FRAP assay of the GFP::GARR-1 transgene in the indicated regions after *rps-8* knockdown. Scale bar, 2 μm. (F) Quantification of FRAP data. Mean ± SD, n ≥ 3 independent animals.

**Figure S3.**
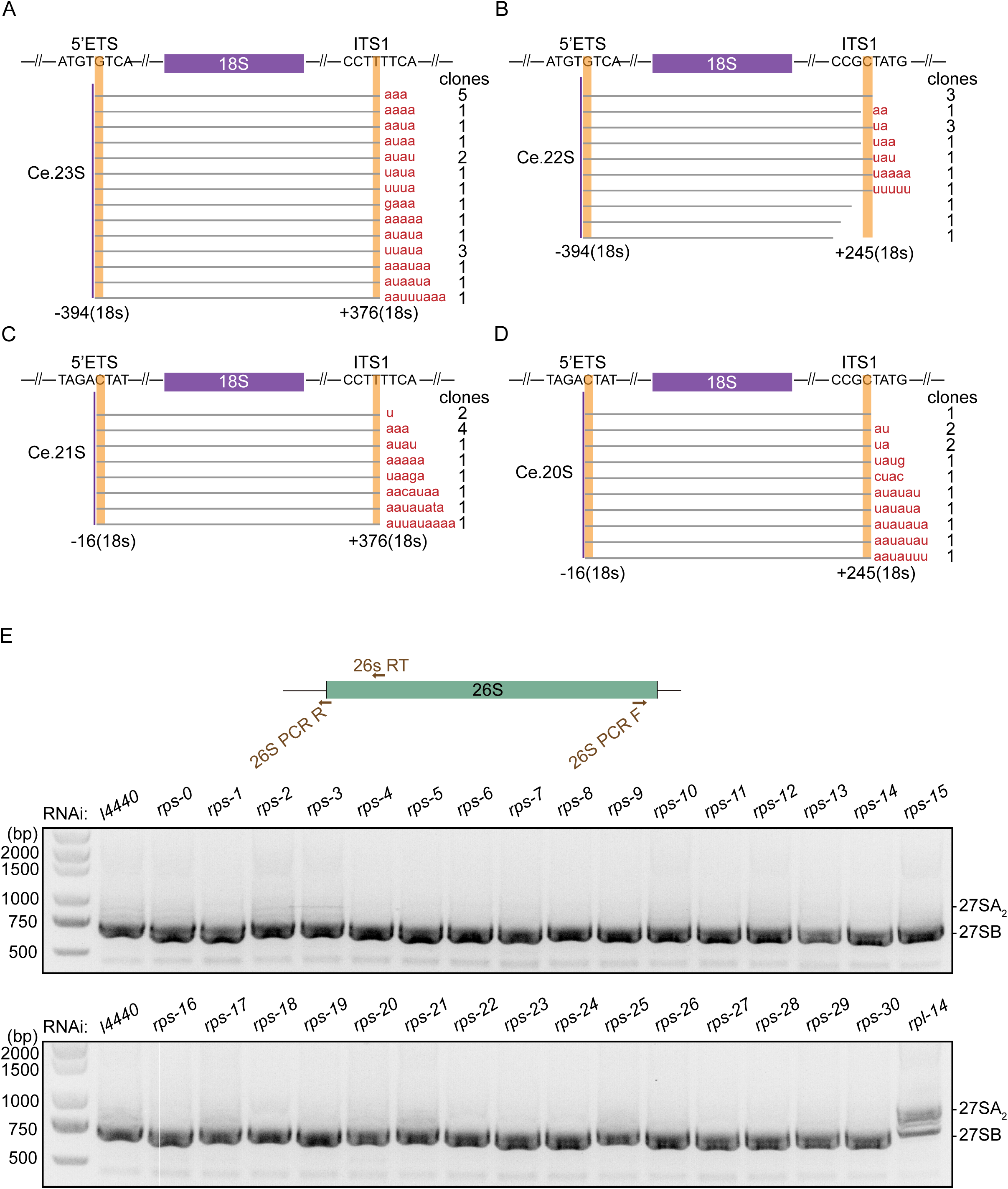
Knocking down class I RPS proteins did not increase the accumulation of 27SA_2_ rRNA intermediate. (A-D) Sanger sequencing of the ends of 18S pre-rRNA intermediates. n represents the number of clones sequenced. (E) cRT‒PCR assay of 27SA_2_ rRNA intermediates upon *rps* knockdown.

**Figure S4.**
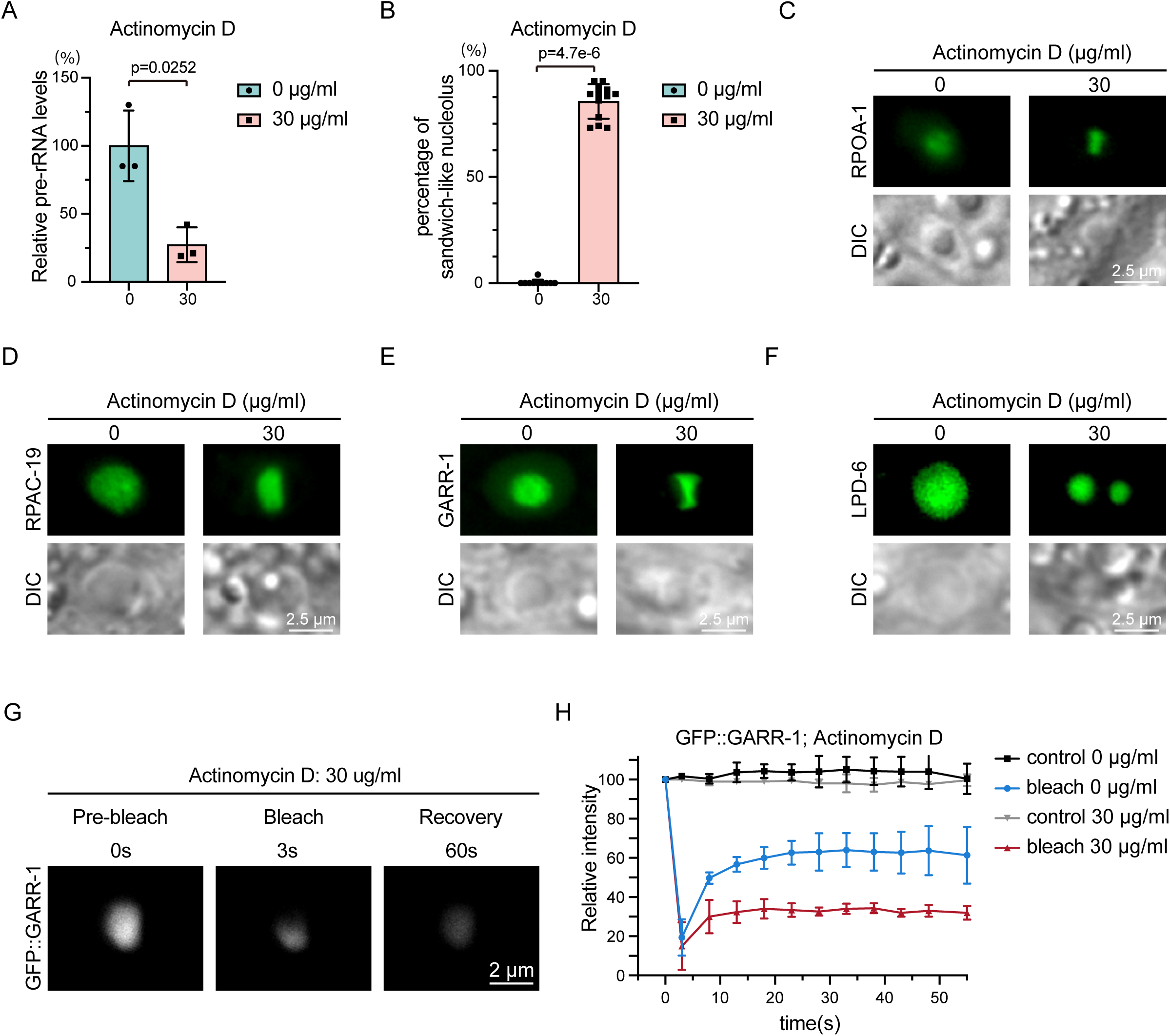
Inhibition of rRNA transcription induces the formation of sandwich-like nucleoli. (A) qRT‒PCR analysis of pre-rRNA levels after 48 h of actinomycin D treatment. The values were normalized to *eft-3* mRNA. The data are presented as the means ± SD of three biological replicates. A two-tailed t test was performed to determine statistical significance. (B) Quantification of the sandwich-like nucleoli after 48 h of actinomycin D treatment. Mean ± SD, n ≥ 10 animals. A two-tailed t test was performed to determine statistical significance. (C-F) DIC and fluorescence microscopy images of the indicated transgenes after actinomycin D treatment. Scale bar, 2.5 μm. (G) FRAP assay of the GFP::GARR-1 transgene in the indicated regions after actinomycin D treatment. Scale bar, 2 μm. (H) Quantification of FRAP data. Mean ± SD, n = 3 independent animals.

**Figure S5.**
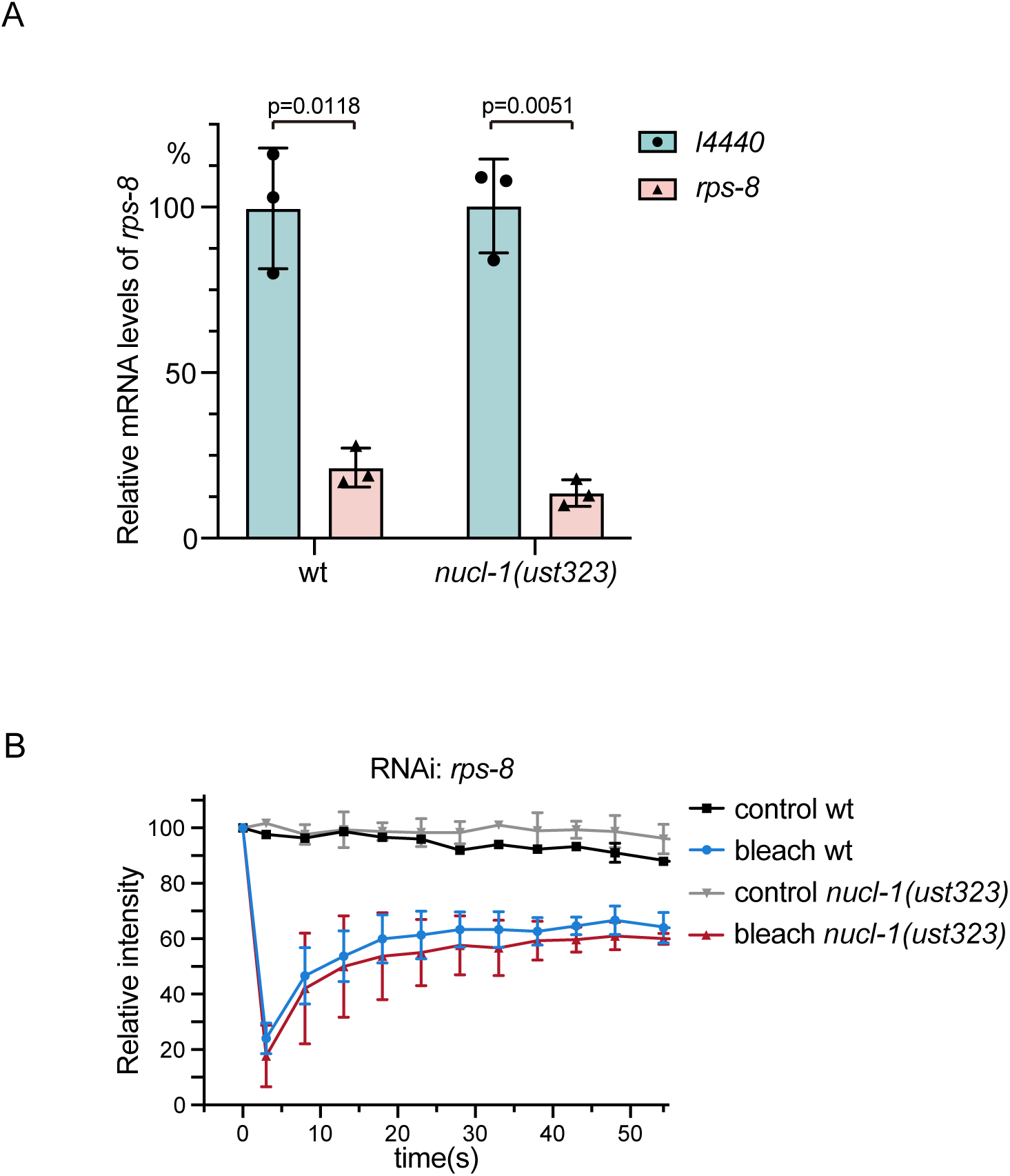
*nucl-1* mutation is not required for nucleolar mobility in *rps-8*-knockdown animals. (A) Relative *rps-8* mRNA levels were measured via quantitative real-time PCR in the indicated animals after *rps-8* RNAi. The data are presented as the means ± SD of three biological replicates. A two-tailed t test was performed to determine statistical significance. (B) FRAP assay of GFP::RPOA-2 in wt and *nucl-1* mutants after *rps-8* knockdown. Mean ± SD, n = 3 independent animals.

**Figure S6.**
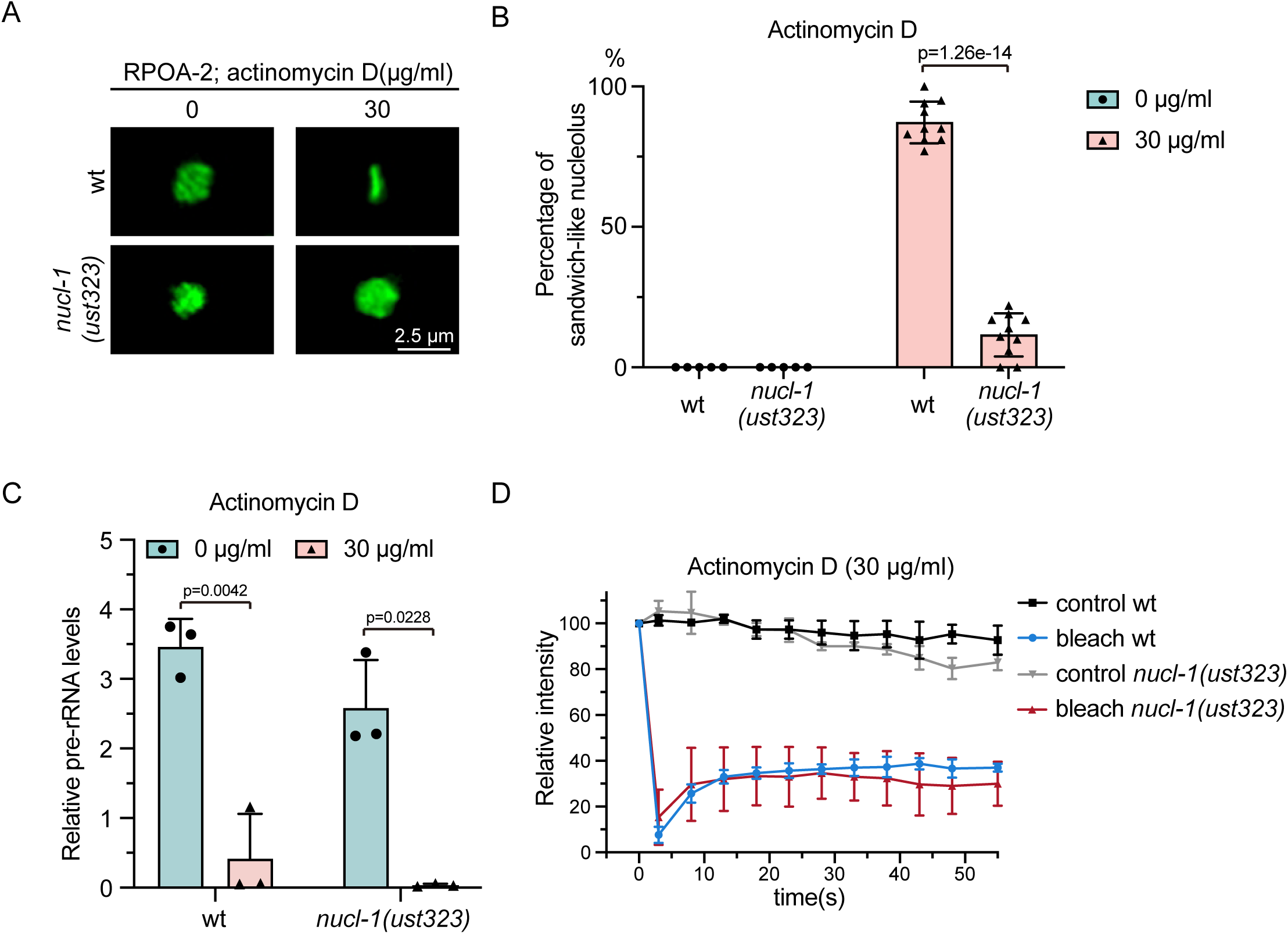
NUCL-1 is required for reorganization of the nucleolar architecture. (A) Fluorescence microscopy images of *C. elegans* nucleoli in hypodermal cells upon actinomycin D treatment in wt and *nucl-1* mutants. Scale bar, 2.5 μm. (B) Quantification of the sandwich-like nucleoli cells upon actinomycin D treatment in wt and *nucl-1* mutants. Mean ± SD, n ≥5 independent animals. A two-tailed t test was performed to determine statistical significance. (C) qRT‒PCR of pre-rRNA levels after the indicated actinomycin D treatment for 48 h in wt and *nucl-1* mutants. The levels were normalized to *eft-3* mRNA. The data are presented as the mean ± SD of three biological replicates. A two-tailed t test was performed to determine statistical significance. (D) FRAP assay of GFP::RPOA-2 in wt and *nucl-1* mutants after actinomycin D treatment. Mean ± SD, n = 3 independent animals.

**Figure S7.**
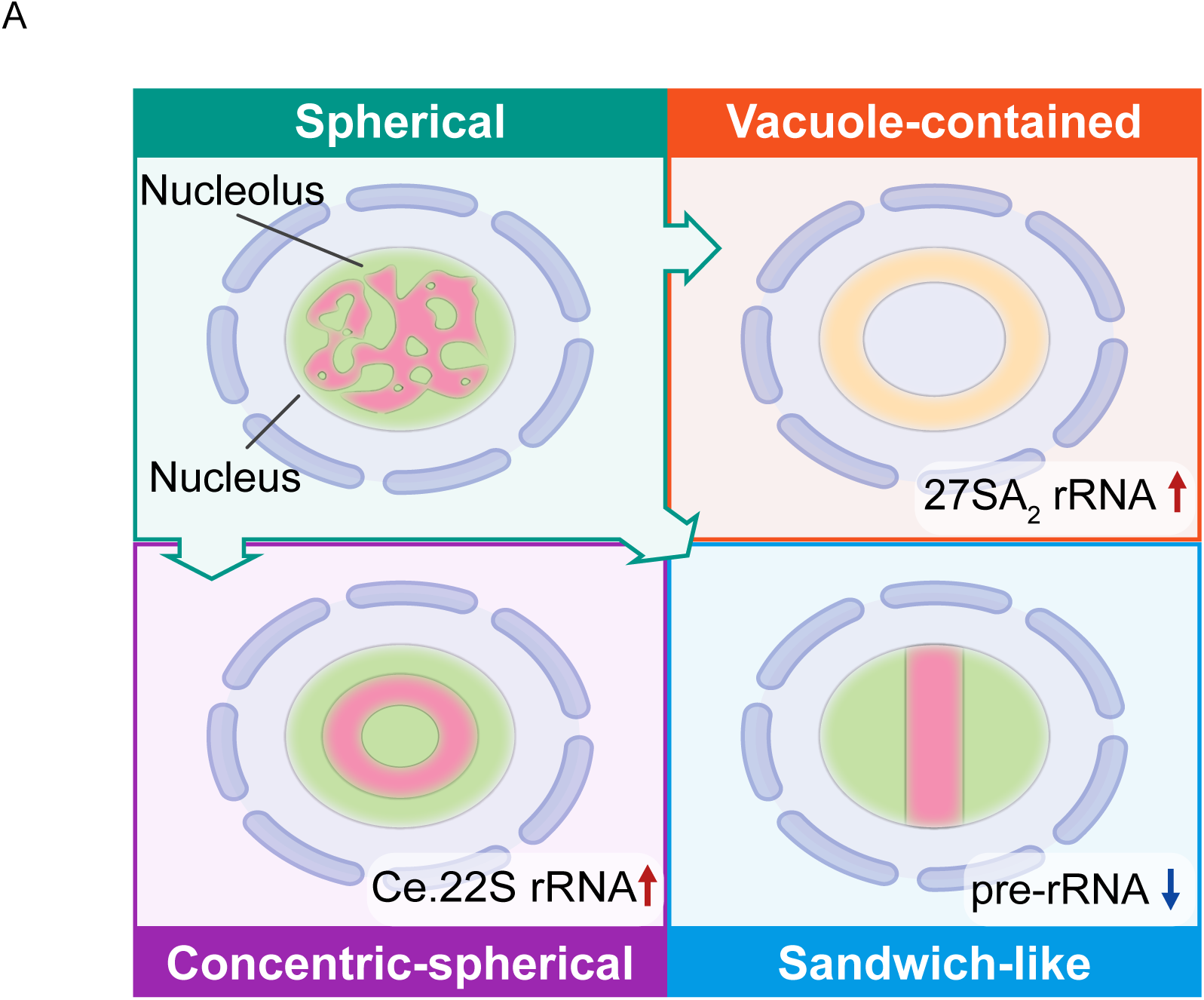
A working model of nucleolar structure reorganization. Diverse impairments in rRNA processing can elicit distinct alterations in nucleolar architecture. The knockdown of the class I *rpl* genes in *C. elegans* leads to inappropriate accumulation of the 27SA_2_ rRNA intermediate, reshapes the nucleolar morphology, and results in the formation of ring-shaped nucleoli and nucleolar vacuoles (NoVs). The accumulation of the Ce.22S rRNA intermediate is induced by knocking down class I *rps* genes, followed by the formation of concentric-spherical nucleoli and demixing of proteins involved in distinct steps of rRNA production. The inhibition of rRNA transcription leads to the formation of sandwich-like nucleoli.

**Table S1.** List of strains used in this study.

| Strain | Genotype |
| --- | --- |
| N2 |  |
| SHG1905 | *rrp-8::gfp; mCherry::rpoa-2* |
| SHG1255 | *eri-1(mg366); mCherry::rpoa-2; 3xflag::gfp::nrde-2* |
| SHG1268 | *3xflag::gfp::rpoa-1 (ustIS137)* |
| SHG1889 | *tagRFP::dao-5 (ustIS242)* |
| SHG1229 | *3xflag::gfp::rpac-19* |
| SHG1660 | *rbd-1::mCherry(ustIS207)* |
| SHG2287 | *fib-1::mCherry* |
| SHG2845 | *gfp::GARR-1(ust579)* |
| SHG2286 | *t06e6.1::mCherry (ustIS292)* |
| SHG2137 | *nucl-1::gfp::3xflag(ustIS279); rrp-8::mCherry* |
| SHG3227 | *lpd-6::gfp* |
| SHG3090 | *3xflag::gfp::rpoa-2 (ustIS116); rps-2p::lacI::tagRFP(ustIS324);*<br>*f31c3.9:: 256xlacO* |
| SHG3104 | *nucl-1::gfp::3xflag(ustIS279); rps-2p::lacI::tagRFP(ustIS324);*<br>*f31c3.9:: 256xlacO* |
| SHG1248 | *3xflag::gfp::rpoa-2 (ustIS116)* |
| SHG2090 | *nucl-1(ust323)* |
| SHG2379 | *nucl-1(ust323); 3xflag::gfp::rpoa-2 (ustIS116)* |

**Table S2.** List of genes used in the RNAi screen.

| Gene ID | Gene ID | Gene ID | Gene ID | Gene ID | Gene ID |
| --- | --- | --- | --- | --- | --- |
| *rps-0* | *rps-1* | *rps-2* | *rps-3* | *rps-4* | *rps-5* |
| *rps-6* | *rps-7* | *rps-8* | *rps-9* | *rps-10* | *rps-11* |
| *rps-12* | *rps-13* | *rps-14* | *rps-15* | *rps-16* | *rps-17* |
| *rps-18* | *rps-19* | *rps-20* | *rps-21* | *rps-22* | *rps-23* |
| *rps-24* | *rps-25* | *rps-26* | *rps-27* | *rps-28* | *rps-29* |
| *rps-29* | *rps-29* | *rps-30* | *rpl-14* |  |  |

**Table S3.**
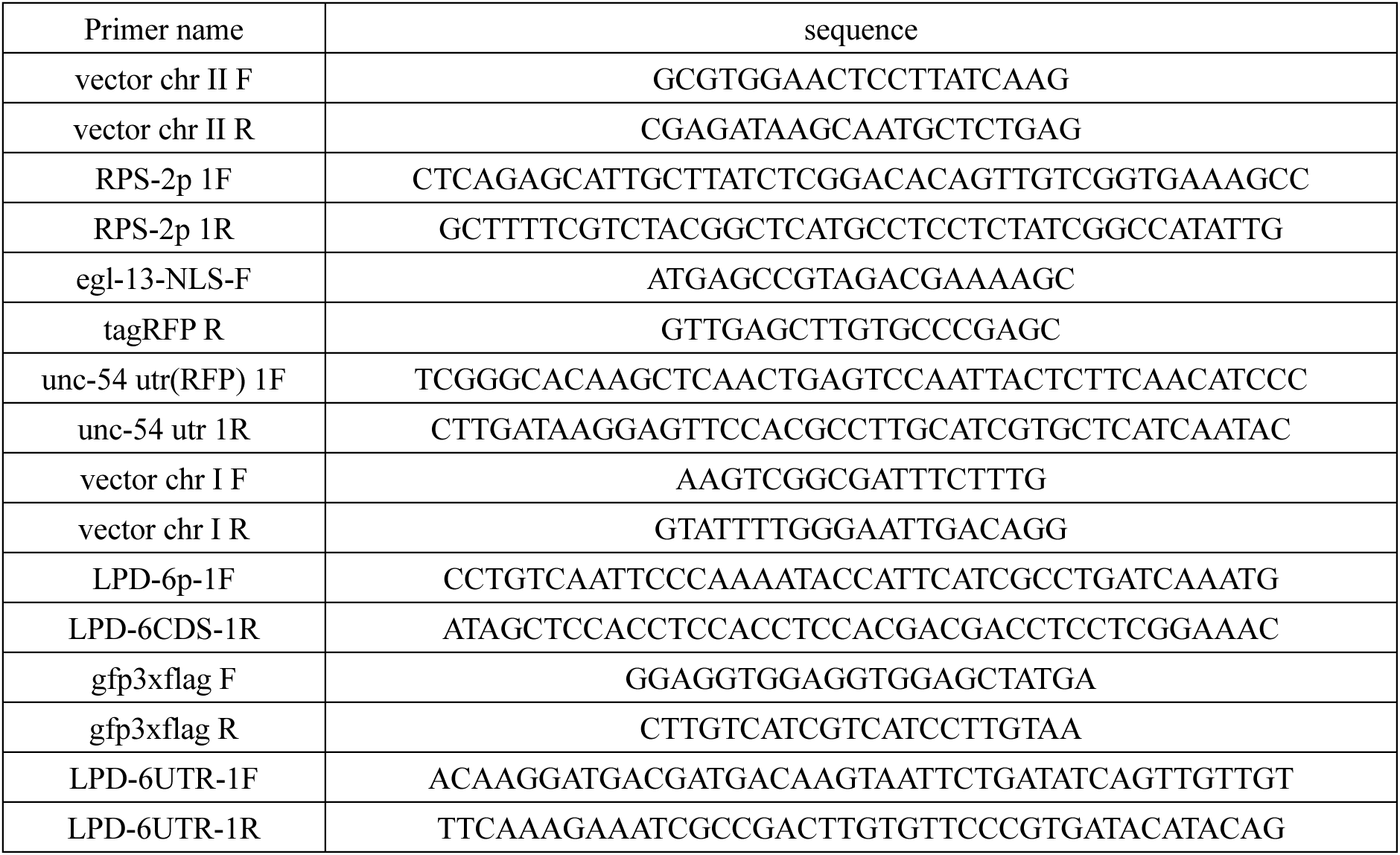
Sequence of repair plasmids primers used in ectopic transgenic strains construction.

**Table S4.**
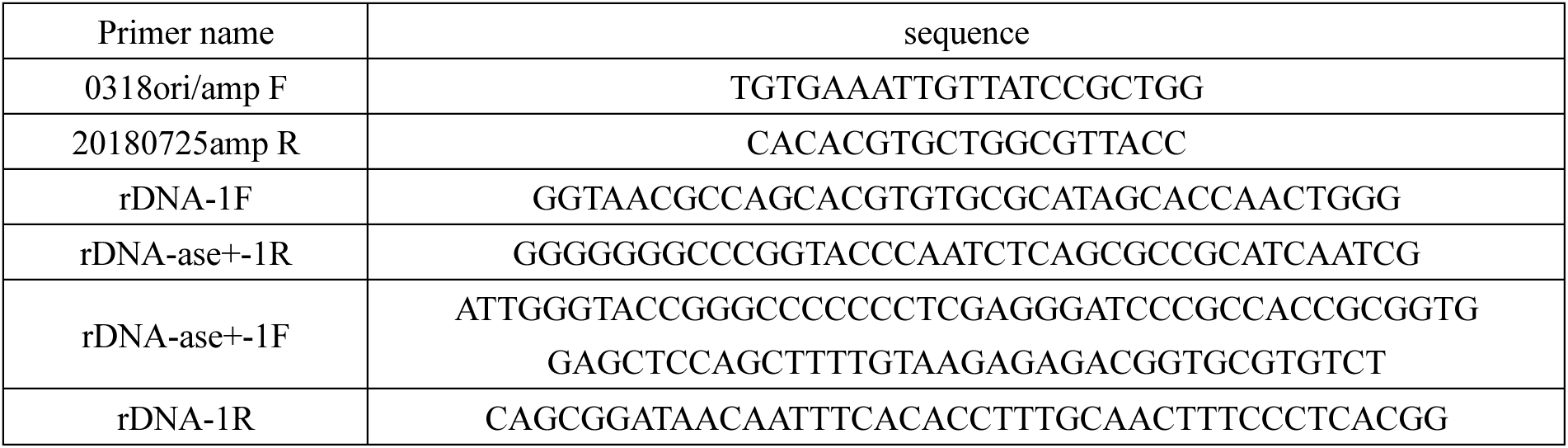
Sequence of repair plasmids primers used in in-situ transgenic strains construction.

**Table S5.**
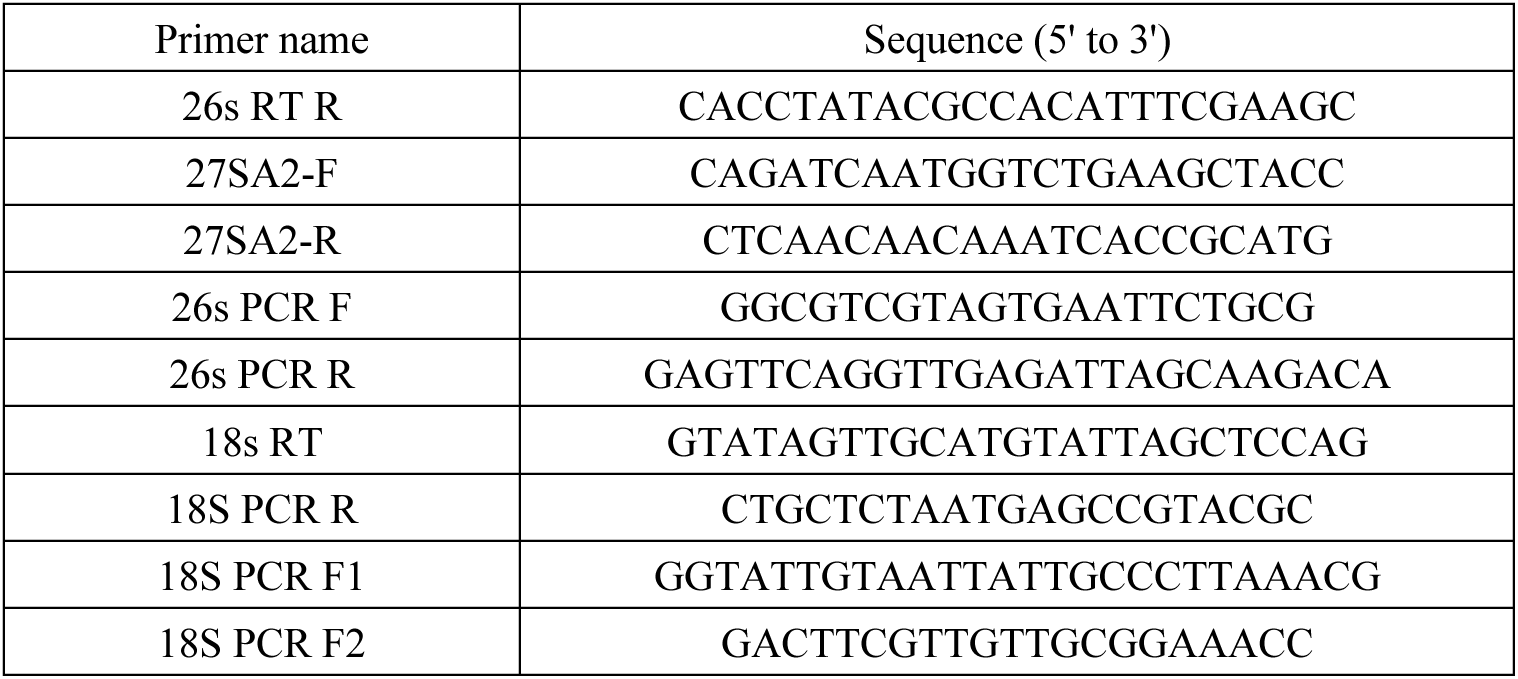
Sequences of cRT-PCR primers.

**Table S6.**
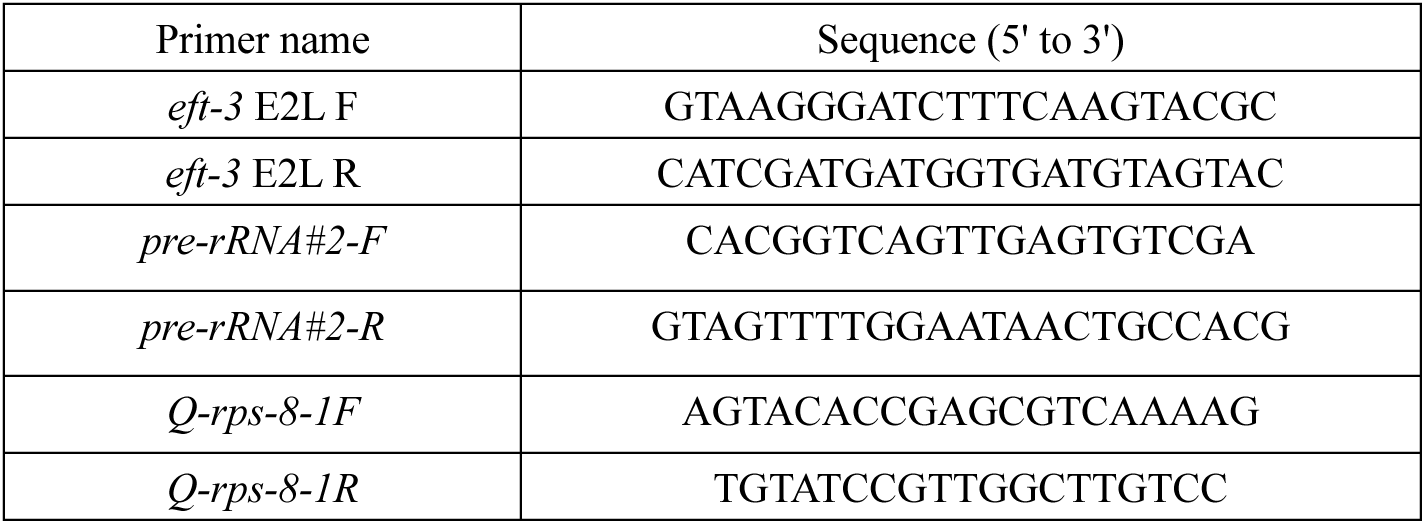
Sequences of qPCR primers.

